# Redox signaling *via* lipid peroxidation regulates retinal progenitor cell differentiation

**DOI:** 10.1101/510438

**Authors:** Shahad Albadri, Federica Naso, Carole Gauron, Carola Parolin, Karine Duroure, Jessica Fiori, Carla Boga, Sophie Vriz, Natalia Calonghi, Filippo Del Bene

## Abstract

Reactive oxygen species (ROS) and downstream products of lipid oxidation are emerging as important secondary messengers in tissue homeostasis. However their regulation and mechanism of action remain poorly studied *in vivo* during normal development. Here we reveal that the fine regulation of hydrogen peroxide (H_2_O_2_) levels at the degradation step by its scavenger Catalase is crucial to mediate the switch from proliferation to differentiation in retinal progenitor cells (RPCs). We further show that altering the levels of downstream products of the Redox signaling can also affect this switch. Indeed, we identify *9*-hydroxystearic acid (*9*-HSA), an endogenous downstream lipid peroxidation product, as a mediator of this effect in the zebrafish retina. In fact, RPCs exposed to higher amounts of *9*-HSA failed to differentiate and remained proliferative. We found that *9*-HSA exerts its biological function *in vivo* by inhibiting the activity of histone deacetylase 1. We finally show that the local and temporal manipulation of H_2_O_2_ levels by *catalase* overexpression in RPCs was sufficient to trigger their premature differentiation. Therefore the amount of H_2_O_2_ in RPCs is instructive of their ability to switch from proliferation to differentiation. We propose a mechanism that acts in RPC and linking H_2_O_2_ homeostasis and neuronal differentiation *via* the modulation of lipid peroxidation.

## Introduction

Adult stem cells and progenitor cells require metabolic plasticity in order to adapt to a quiescent or highly proliferative state, respectively. Stem cells preference for aerobic glycolysis rather than oxidative phosphorylation has been proposed to be due to the hypoxic environment in which they replicate and the relatively low energy requirement because of their quiescent status (Bigarella *et al*, 2014; Norddahl *et al*, 2011; Yeo *et al*, 2013). Already a decade ago, Prozorovski et al. have highlighted the role of the redox signaling in the regulation of neural progenitor fate (Prozorovski *et al*, 2008). Thus, the redox pathway has been proposed to regulate neural stem cell self-renewal and neurogenesis *via* PI3K/Akt signaling, further highlighting the physiological need of ROS molecules as cellular signaling molecules (Le Belle *et al*, 2011). Other studies have shown that tumor cells can also highjack the redox pathway by modulating their endogenous levels of ROS and downstream products for the tumor progression to occur (Gorrini *et al*, 2013; Piskounova *et al*, 2015; Zhou *et al*, 2014). Overall all these reports highlight the importance of ROS level regulation for them to achieve their physiological function. However to this date, the *“metabolic master switch that determine the fate of neural progenitors*” remain uncharacterized (Prozorovski *et al*, 2008).

In the last years, H_2_O_2_ has emerged as the major redox metabolite that acts as an essential messenger for several fundamental processes such as inflammation, hypoxia, regeneration and wound-healing but also proliferation and stem cell self-renewal, tumorigenesis and aging, diabetes, etc. (Holmström & Finkel, 2014; Sies *et al*, 2017; Sies, 2017). In fish and amphibians, cell proliferation subsequent to injury was shown to correlate with the increase in the amounts of H_2_O_2_ at the site of amputation (Niethammer *et al*, 2009; Love *et al*, 2013; Hameed *et al*, 2015). During the central nervous system (CNS) development, we and others have reported a physiological function for H_2_O_2_ in controlling axon growth and regeneration (Gauron *et al*, 2016; Meda *et al*, 2016; Hervera *et al*, 2018; Rieger & Sagasti, 2011; Wilson *et al*, 2017). In our previous study, we revealed the dynamic landscape of the molecule during morphogenesis *in vivo*. We were able to demonstrate that the levels of H_2_O_2_ during this process were regulated at the degradation step to ensure their presence at optimal levels (Gauron *et al*, 2016).

In the present study, we sought to determine the role of the redox pathway and lipid peroxidation products as its possible mediators during the development of the zebrafish neuroretina, taking advantage of its well-defined architecture and cell organization. At 2 days post fertilization (dpf) during zebrafish retinogenesis, retinal progenitor cells (RPCs) start differentiating in a central to peripheral manner. By 3 dpf, the central retina is fully differentiated and a peripheral stem cell niche, known as the ciliary marginal zone (CMZ), forms to ensure tissue homeostasis throughout the lifelong growth of the fish. The CMZ comprises three main cell domains labeled by the expression of different transcription factors: the *rx2* domain, which contains RSCs (retinal stem cells) and RPCs, the *ccnd1* (zebrafish equivalent to *cyclinDl)* domain where cycling progenitors are found and an *ath5* domain, which contains cycling and committed RPCs (Agathocleous & Harris, 2009; Ohnuma & Harris, 2003; Reinhardt *et al*, 2015; Wan *et al*, 2016). Due to this precise spatial and temporal organization, the zebrafish retina provides an excellent model to assess the effects of ROS and lipid peroxidation downstream products on a stem cell and progenitor cell population *in vivo.* Here we show that H_2_O_2_ levels are dynamically regulated in the developing and in the mature neuro-retina where the expression of the H_2_O_2_ scavenger *catalase* follows a pattern reminiscent of the neurogenic progression. H_2_O_2_ levels are further reflected on the lipid peroxidation state in these cells as shown by the enrichment in the end marker of lipid peroxidation 4-hydroxynonenal (4-HNE) within the CMZ. Thus, H_2_O_2_ and lipid peroxidation may be act in the regulation of retinal cell proliferation and differentiation during the development and the growth of the tissue.

We previously characterized the *in vitro* role of the *9*-hydroxystearic acid (*9*-HSA), which was found endogenously produced in human embryonic intestine cells and colon adenocarcinoma as signaling messenger produced by lipid peroxidation (Piretti *et al*, 1987; Piretti & Pagliuca, 1989). *9*-HSA amounts in cancer cells were found inversely correlating with tumor growth (Bertucci *et al*, 2002; Calonghi *et al*, 2007; Cavalli *et al*, 1991; Gesmundo *et al*, 1994). We found that in zebrafish, *9*-HSA is endogenously produced and its exogenous administration to RPCs severely altered their differentiation, identifying it as a critical molecule acting downstream of redox signaling events.

Our study reveals the physiological role of the redox pathway and lipid peroxidation *in vivo* during retinal development and for the homeostasis maintenance of the mature retina. Our results bring to light the cascade of events that leads to the up-regulation of pro-neuronal genes in RPCs, which are required for transition from proliferation to differentiation.

## Results

### ROS scavenging and lipid peroxidation are active during retinal development

ROS homeostasis, resulting from their production and degradation within a tissue, is tightly regulated as shown for H_2_O_2_ regulation during morphogenesis (Gauron *et al*, 2016). With the aim of uncovering the dynamic of H_2_O_2_ production in the retina during its development, we first used the ratiometric sensor transgenic line Tg(*ubi:HyPer*) similar to one we previously established as a reporter for the endogenous levels of H_2_O_2_ (Gauron *et al*, 2016). We imaged zebrafish retinae at two specific developmental stages: at 24 hours post-fertilization (hpf) when the undifferentiated neuroepithelium is highly proliferative and at 32 hpf when differentiation has started in the central part of the tissue (Figure 1a – a’). We could observe a strong correlation between cell proliferation and high levels of H_2_O_2_. Indeed at 24 hpf, high amounts of H_2_O_2_ could be observed throughout the epithelium whereas at 32 hpf, the central part of the tissue was depleted of H_2_O_2_ (Figure 1a – a’). Indeed, the measurement of H_2_O_2_ levels across the tissue showed that H_2_O_2_ levels remain high at the periphery of the developing retina where retinal progenitors are still proliferating, while they drastically decreased in the central part of the tissue where differentiation has been initiated (Figure 1b). These results reveal the dynamics of H_2_O_2_ production in relation to cell differentiation and proliferation in the retina and suggest a putative role of this pathway for the development of the tissue.

**Figure 1.**
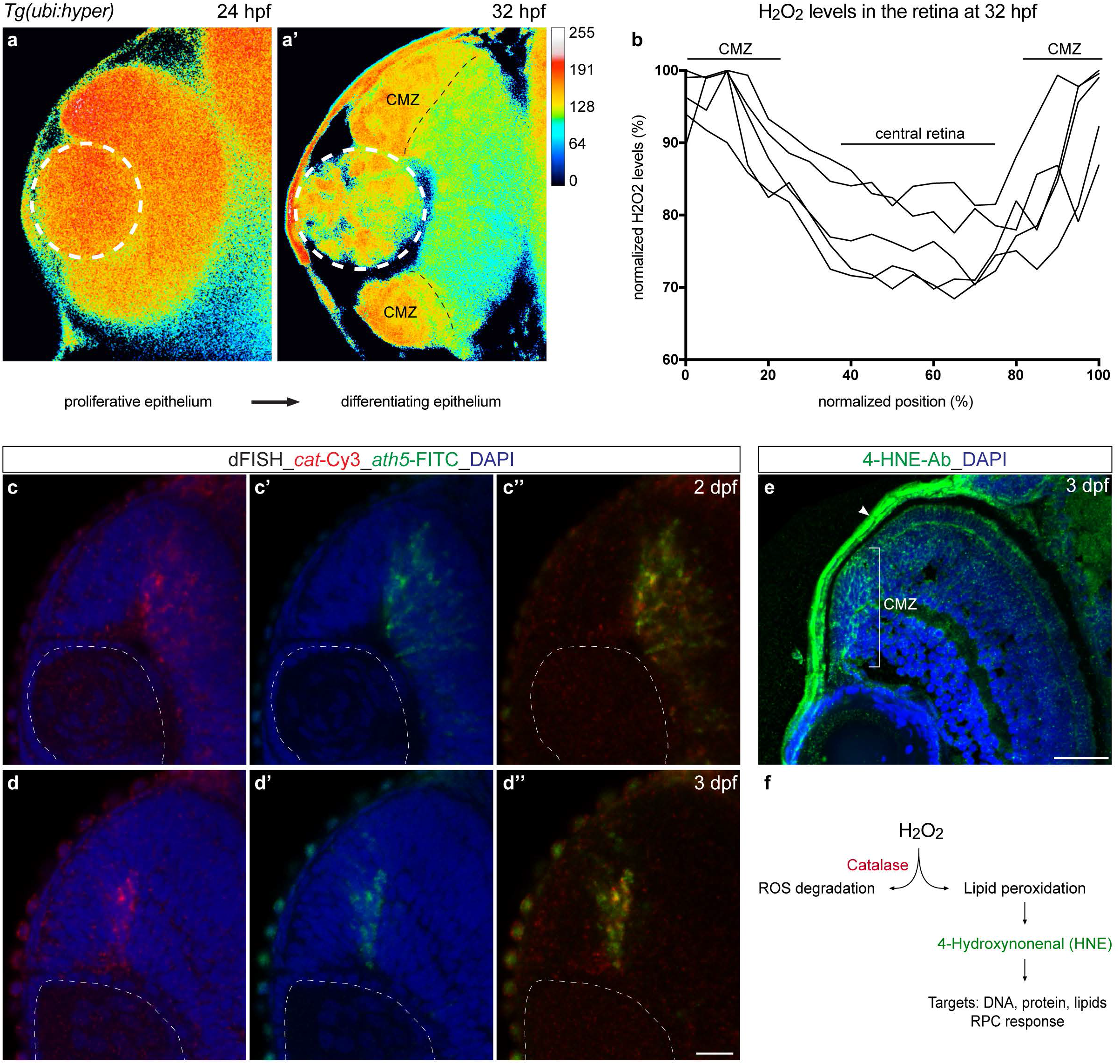
H_2_O_2_ production, scavenging and lipid peroxidation processes are active in the zebrafish retina. (a – b) H_2_O_2_ levels change in relation to retinal development revealed by the reporter line Tg(*ubi:HyPer*). The H_2_O_2_ levels are inferred from the YFP_500_/YFP_420_ excitation ratio of HyPer. (a – a’) Optical section of a Tg(*ubi:HyPer*) retina at 24 hpf shows homogeneous high levels of H_2_O_2_ throughout the proliferative epithelium. At 32 hpf, H_2_O_2_ content decreased in the central part of the tissue and remained high in the ciliary marginal zone (CMZ) as confirmed by normalized quantification of H_2_O_2_ levels at this stage (b). *Catalase* was expressed in cycling retinal progenitor cells independently from Ath5 function. (c – d’’) *Catalase* and *ath5* transcripts overlap during retinogenesis at 2 and 3 days post-fertilization (dpf). Confocal sections through the central retina of wild type embryos hybridized with both *catalase* (red) and *ath5* (green) antisense RNA probes and counterstained with the nuclear marker DAPI (blue). At 2 dpf, both genes are expressed in the central retina transiently (c – c’’) and are confined to the Ath5 domain of the ciliary marginal zone at 3 dpf (d – d’’). White dashed lines delimitate the lens. Scale bar, 20 μm. The lipid peroxidation end marker 4-HNE is endogenously produced in zebrafish. (e) Lipid peroxidation marker 4-HNE was enriched in the ciliary marginal zone of the 3 dpf zebrafish retina. Immunohistochemistry of 4-HNE (green) on 3 dpf frontal retinal cryosection counterstained with DAPI (blue) for nuclear labeling. The white arrow points at 4-HNE staining at the surface of the skin where high oxidative stress processing occurs as expected. Scale bar, 100 μm. (f) Schematic representation of H_2_O_2_ scavenging *via* Catalase coupled with the lipid peroxidation pathway and the downstream production of 4-HNE as secondary messenger.

We secondly aimed at assessing how H_2_O_2_ scavenging occurs in the zebrafish retina over time. We first evaluated the expression dynamics of *catalase*, which encodes for the antioxidant enzyme Catalase. Catalase, which transforms H_2_O_2_ into water and oxygen, is at the end of the antioxidant pathway that regulates cellular oxidative state oscillations, providing a protection against ROS-induced damages (Lynch & Fridovich, 1978; Rhee *et al*, 2005; Sies, 1991; Spitz *et al*, 2004). We previously reported that *catalase* expression occurs mainly in the brain where fine-tuning of H_2_O_2_ levels is achieved by degradation (Gauron et al, 2016). By double fluorescent *in situ* hybridization experiments at 2 and 3 days post-fertilization (dpf), we were able to detect *catalase* mRNA and to relate it to the expression of the bHLH transcription factor *ath5*, a marker for retinal neurogenesis initiation and later for cycling retinal progenitor cells (RPCs) in the ciliary marginal zone (CMZ) (Fig. 1c – c’’) (Kay *et al*, 2005). *Ath5* and *catalase* transcripts were found to co-localize at 2 dpf in the central retina where they are transiently expressed (Fig. 1c – c’’) and at 3 dpf in the CMZ (Fig. 1d – d’’). In order to assess the relationship between the two genes, we next monitored *catalase* expression in the *ath5-^-/-^* mutant, also known as *lakritz* (Supplementary Fig. 1A-B) (Kay *et al*, 2001). Interestingly, the absence of Ath5 in the retina did not affect *catalase* expression, which is still found in the CMZ (Supplementary Fig. 1A – B). Therefore *catalase* is expressed in the same neurogenic domain labeled by *ath5*, but independently from Ath5 function itself. These results indicate that *catalase* is expressed in cycling RPCs prior to their last division during development and, later, during retinal growth.

ROS including H_2_O_2_ can react with polyunsaturated fatty acids of the lipid membrane and induce lipid peroxidation. 4-hydroxynonenal (4-HNE) is a known end-product of lipid peroxidation and is considered as a secondary messenger and marker of oxidative stress (Barrera *et al*, 2008; Barrera, 2012; Pizzimenti *et al*, 2010). In order to ask whether lipid peroxidation occurs in RPCs and differentiated neurons, we assessed their content in 4-HNE at 3 dpf (Schneider *et al*, 2001). By immunohistochemistry, we were able to detect enrichment in 4-HNE signals in the CMZ, while little or no signal was detectable in the differentiated part of the 3 dpf retina (Fig. 1e). These results reveal that, in contrast to differentiated retinal neurons, RPCs have a higher content in 4-HNE, indicating that lipid peroxidation is active within these cells.

Taken together, our results indicate that H_2_O_2_ scavenging and lipid peroxidation are active processes occurring within cycling RPCs during retinogenesis and retinal growth (Fig. 1f).

### *9*-HSA lipid peroxidation product is produced endogenously in zebrafish

In the search of a possible molecular mechanism that could link H_2_O_2_ homeostasis and lipid peroxidation to cell proliferation and differentiation in the retina, we focused our interest on the *9*-hydroxystearic acid (*9*-HSA) as one of the putative endogenous mediator of the redox pathway in the retina. *9*-HSA has indeed previously been identified as an endogenous by-product of the lipid peroxidation in colon carcinoma cells where it acts as a growth inhibitor (Cavalli *et al*, 1991; Calonghi *et al*, 2007; Gesmundo *et al*, 1994). Given the importance in human cancer cell models, we assessed whether *9*-HSA is endogenously produced *in vivo* in zebrafish under physiological conditions. In order to evaluate the content of *9*-HSA in the lipid extract of the zebrafish embryo, a liquid chromatography-electrospray ionization-tandem mass spectrometry (LC-ESI-MS/MS) method was optimized. The reverse phase chromatographic separation efficiency was evaluated by analyzing simultaneously *9*-HSA and its isomer *10*-HSA and, as it can be observed in the chromatogram in Fig. 2a, the analysis allowed a good separation of the two molecules. A calibration curve (y = 283,78x – 42295) was obtained by plotting the fragment ion 253.2 peak area *versus* the corresponding concentration, expressed as ng/mL. The linear correlation coefficient (r2) exceeded 0.995, indicating a linear behavior of the response over two orders of concentration of the analytes. Quantification of the endogenous amount of *9*-HSA in zebrafish embryos at different developmental stages (prior 1 dpf, 1, 3 and 4 dpf) was performed and results were expressed as *μ*g/mg of lipid extract. We observed an increase in the content of *9*-HSA in the lipid extract of the zebrafish embryo over time (Fig. 2b). Our quantifications therefore reveal that *9*-HSA is endogenously produced during zebrafish development. Taken together, the biological role of *9*-HSA in cell culture and its endogenous production by the lipid peroxidation machinery in zebrafish suggest an *in vivo* role for the molecule (Fig. 2c).

**Figure 2.**
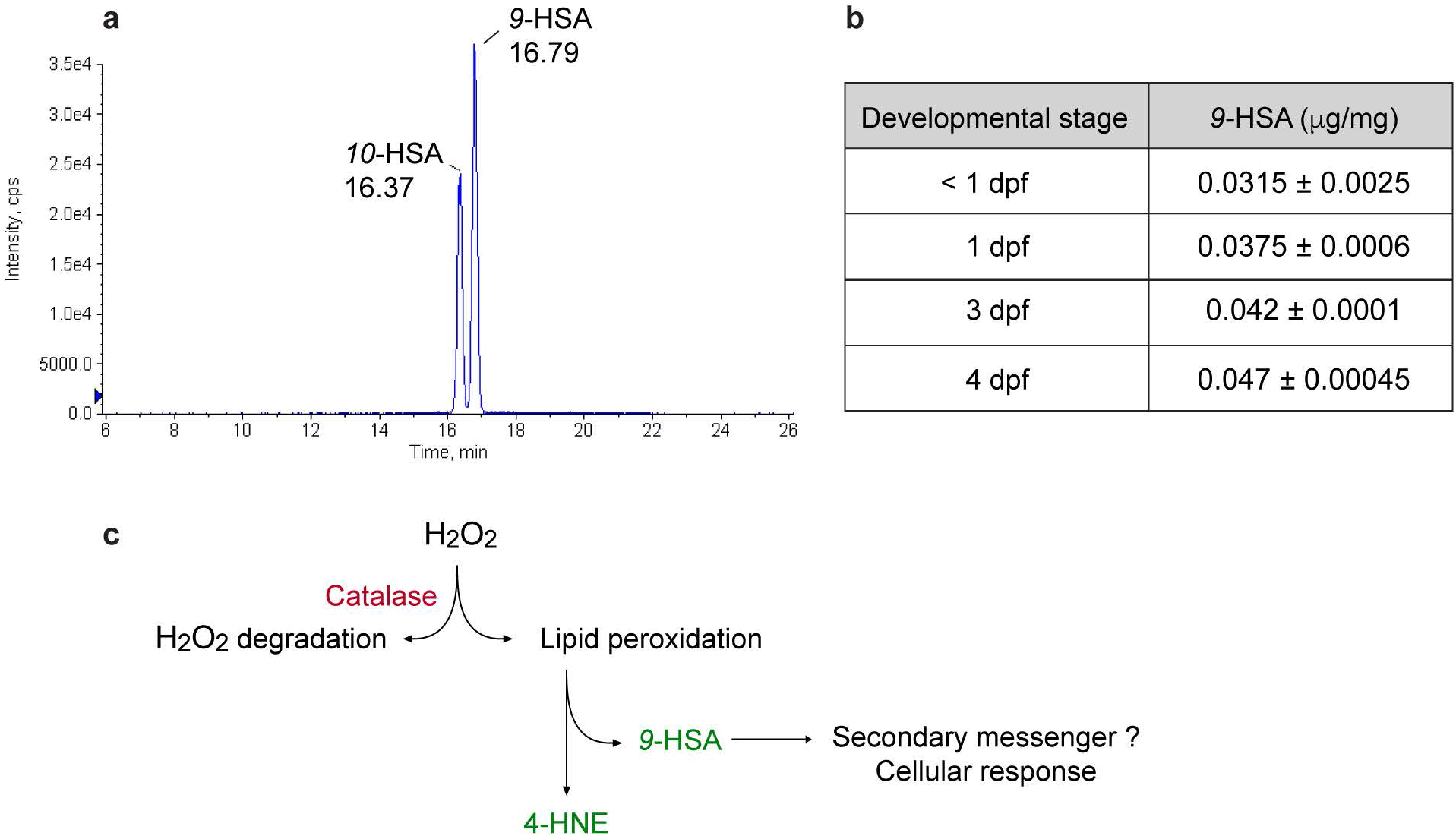
*9*-HSA lipid peroxidation product is endogenously produced in zebrafish. (a) Liquid chromatography tandem mass spectrometry (LC-ESI-MS/MS) chromatogram of *9*-HSA and *10-* HSA. The reverse phase chromatographic separation efficiency was evaluated by analyzing simultaneously *9*-HSA and its isomer 10-HSA. The analysis allowed a good separation of the two molecules. (b) Quantification of *9*-HSA endogenous amounts in zebrafish embryos of different developmental stages: prior to 1, 1, 3 and 4 dpf. An increase in the content of *9*-HSA during development was observed. Extraction results are expressed in μg/mg. (c) Schematic representation of ROS scavenging *via* Catalase coupled with the lipid peroxidation pathway and the downstream production of *9*-HSA as secondary messenger.

### Exogenous administration of *9*-HSA leads to retinal differentiation defects

We next asked whether *9*-HSA has a role in the regulation of RPC fate. To do so, we evaluated the effect of RPC overexposure to *9*-HSA by exogenous administration on retinal differentiation. Synthetic *9*-HSA (or DMSO vehicle) was injected into 1-cell stage zebrafish embryos and retinal sections were analyzed at 2 and 3 dpf (Fig. 3). At 2 dpf, retinal ganglions cells (RGCs) have differentiated, and at 3 dpf the fish retina is fully differentiated and comprises all neuronal and glia cell types (Hu & Easter, 1999). Immunohistochemistry for different retinal cell classes was performed: Zn5 marking all retinal ganglion cells at 2 dpf, Parvalbumin for amacrine cells, GS for Müller glia cells and Zpr1 for photoreceptors at 3 dpf (respectively Fig. 3a, b, c, d). With respect to the DMSO control retinae, our analysis revealed a strong reduction in differentiation markers expression for all the tested retinal cell types in the *9*-HSA injected embryos (Fig. 3a’, b’, c’, d’ respectively). In some cases, like for Zpr1 photoreceptor marker, the most central part of the retina appeared to be more affected than the peripheral regions (Fig. 3d’). This can be explained by the retinal differentiation process, which temporally propagates from the central retina to the periphery (Stenkamp, 2007). Therefore cells in the distal part undergo neuronal differentiation at later developmental stages, when exogenous *9*-HSA amounts may have started to decrease. These observed defects in differentiation in *9*-HSA –treated retinae were not associated with cell death as can be observed at 3 dpf by TUNEL assay (Supplementary Fig. 2). Together our results demonstrate that RPCs overexposed to *9*-HSA during early development fail to differentiate without however undergoing apoptosis at the assessed developmental stage.

**Figure 3.**
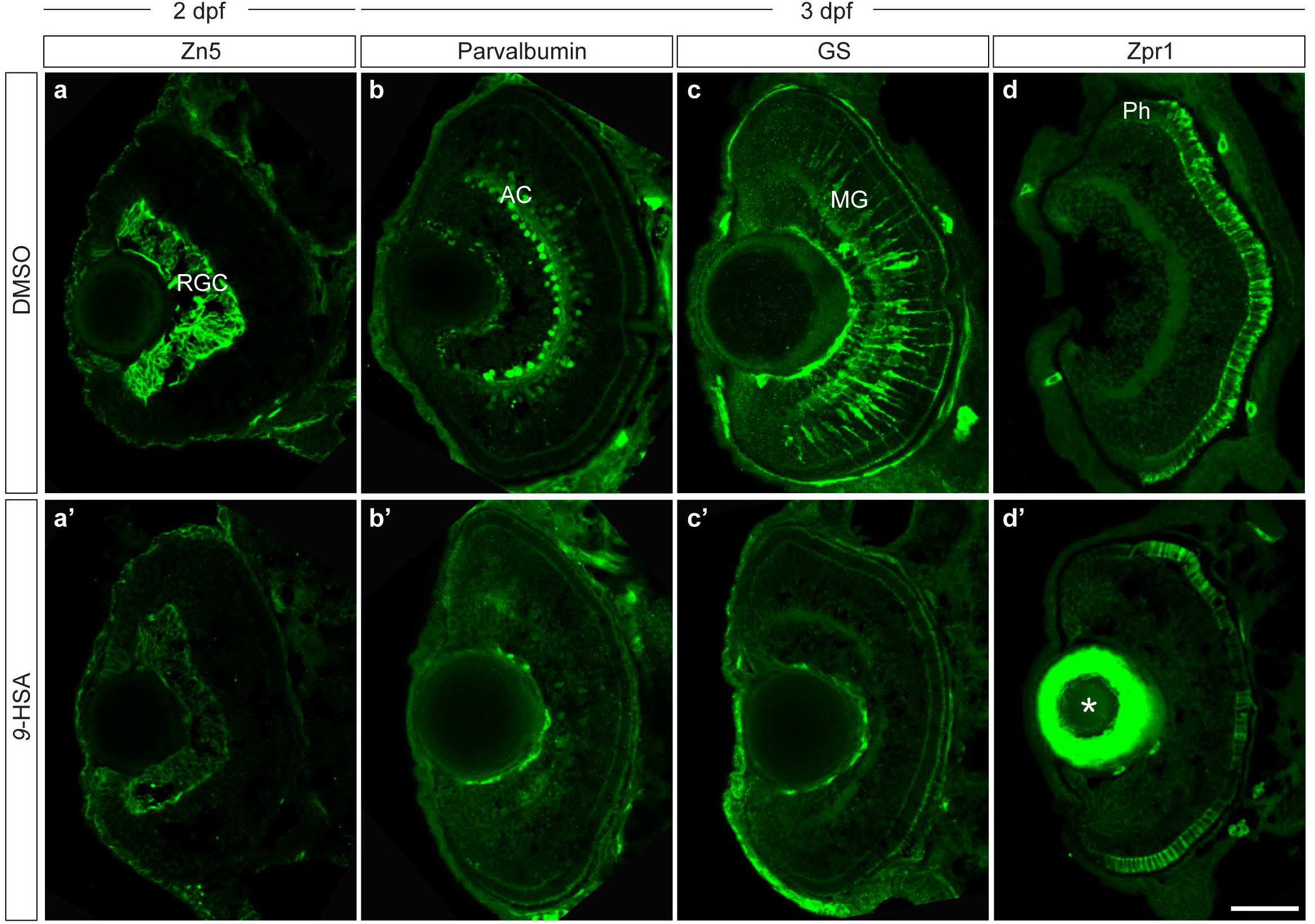
Overexposure of retinal progenitor cells to *9*-HSA leads to defects in differentiation. (a – d’) *9*-HSA exogenous administration leads to defects in differentiation during retinal development for all cell types analyzed. (a – a’) Immunohistochemistry of anti-Zn5 (green) labeling retinal ganglion cells (RGC) on 2 days post-fertilization (dpf) frontal retinal cryosections from DMSO-treated (a) and *9*-HSA -treated (a’) embryos. No RGC labeling was observed in the central part of the retina at 2 dpf in *9*-HSA -treated in comparison to DMSO-treated control retinae. (b – b’) Immunohistochemistry of anti-Parvalbumin (green) labeling amacrine cells and displaced amacrine cells (AC) on 3 dpf frontal retinal cryosections from DMSO-treated (b) and *9*-HSA -treated (b’) embryos. Similarly to RGCs, no amacrine cells or displaced amacrine cells were observed in *9*-HSA -treated in comparison to DMSO-treated control retinae. (c – c’) Immunohistochemistry of anti-GS (green) labeling Müller glia cells (MG) on 3 dpf frontal retinal cryosections from DMSO-treated (c) and *9*-HSA –treated (c’) embryos revealed no MG differentiation in *9*-HSA -treated in comparison to the DMSO control retinae. (d – d’) Immunohistochemistry of anti-Zpr1 (green) labeling photoreceptor cells (Ph) on 3 dpf frontal retinal cryosections from DMSO-treated (d) and *9*-HSA -treated (d’) embryos. No Zpr1 labeling was observed in the central part of *9*-HSA -treated retinae compared to DMSO-treated control retinae where Zpr1-positive Ph could be detected within the outer nuclear layer of the 3 dpf retinal tissue. Asterisk in (d’) indicates unspecific antibody trapping in the lens. Scale bar, 50 μm.

### RPCs exposed to high levels of *9*-HSA remain proliferative

*9*-HSA supplemented in micro-molar amounts to the human colon cancer cell line HT29 was previously shown to result in a strong inhibition of cell proliferation as well as cellular differentiation toward benign phenotype (Calonghi *et al*, 2005). To evaluate the effect of *9*-HSA on the proliferation of RPCs during retinogenesis and later on the stem cell niche of the retina, we assessed the proliferative state of retinae from embryos injected with *9*-HSA. Under physiological conditions, at 2 dpf the retinal neuroepithelium is mainly differentiated in the central part and by 3 dpf RPCs cells are restricted to the CMZ domain while the central part of the retina is entirely occupied by post-mitotic neurons and glia cells. A 12 hours BrdU incorporation assay labeling cells that entered the S-phase was performed on 2 dpf *9*-HSA -injected and DMSO-injected embryos (Fig. 4a – a’). Compared to DMSO control retinae where BrdU could be detected mostly in the periphery of the tissue in the nascent CMZ at 2.5 dpf, we could detect BrdU incorporation in the central part of the retina in *9*-HSA -treated embryos (Fig. 4a – a’). Immunohistochemistry performed with antibodies against the proliferating cell nuclear antigen (PCNA, S-phase marker) and against the phosphorylated histone H3 (pH3, M-phase marker) was carried out on 2 and 3 dpf retinal sections from *9*-HSA -injected and DMSO-injected embryos (Fig. 4b – d). *9*-HSA-treated retinae showed an expanded PCNA-positive CMZ region and PCNA labeling in the central retina compared to the control situation (Fig. 4b - b’) and the proportion of pH3-positive cells in *9*-HSA retinae were significantly increased at 2 and 3 dpf (n = 5, p-value: 0,0176 at 2 dpf and n = 4, p-value: 0,0060 at 3 dpf for *9*-HSA, Fig. 4c - d). These results indicate that RPCs overexposed to *9*-HSA amounts fail at exiting the cell cycle and are more prone to remain in their proliferative state, in contrast to *9*-HSA role *in vitro* where it acts as a growth inhibitor.

**Figure 4.**
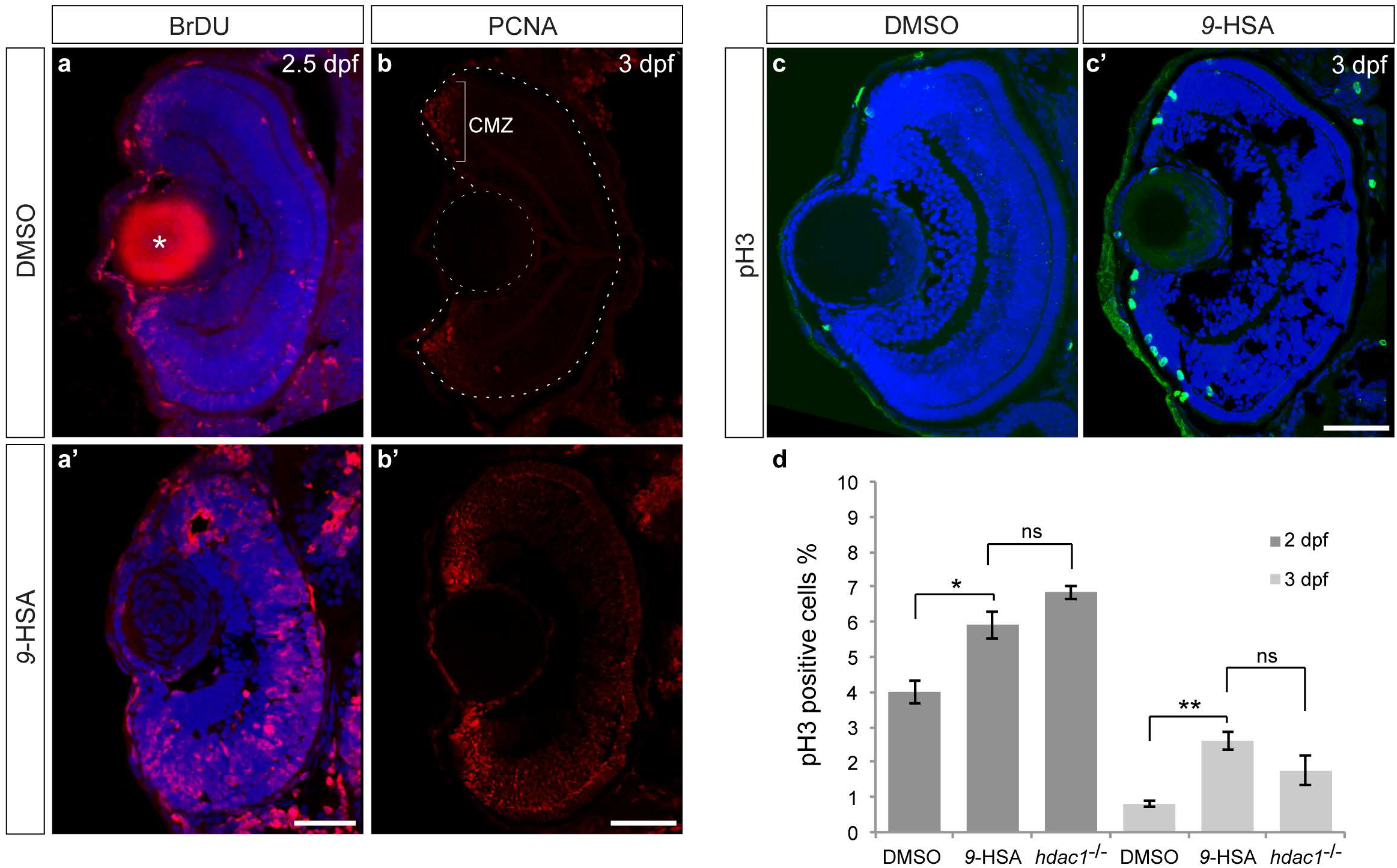
*9*-HSA exogenous administration leads to an increase in proliferation in the retina. (a – a’) BrdU incorporation assay at 2 days post-fertilization (dpf) in DMSO-treated control and *9*-HSA – treated embryos detected by immunohistochemistry after 12 hours shows a strong increase in the number of cells that actively replicated their DNA (S-phase, red) during these 12 hours. Nuclear counterstaining was also performed on these retinal sections by DAPI labeling (blue). (b – b’) Immunohistochemistry of anti-PCNA (red) labeling cells in S-phase of the cell cycle on 3 dpf frontal retinal cryosections. *9*-HSA –treated retinae showed an expanded PCNA labeling in the ciliary marginal zone (CMZ, bracket) and in the central retina in comparison to DMSO –treated control retinae at 3 dpf. (c – c’) Immunohistochemistry of anti-pH3 (green) labeling cells in M-phase of the cell cycle on 3 dpf frontal retinal cryosections counterstained with the nuclear marker DAPI (blue). *9*-HSA –treated retinae showed a higher number of mitotic cells in comparison to DMSO –treated control retinae at 3 dpf. (d) Quantification of the mitotic index at 2 and 3 dpf shows a significant increase in the percentage of pH3-positive cells in *9*-HSA –treated retinae with respect to DMSO – treated control retinae at 2 dpf and 3 dpf (n = 5, p-value: 0.0176 at 2 dpf and n = 4, p-value: 0.0060 at 3 dpf). No significant difference in the mitotic index was observed between *9*-HSA –treated retinae and *hdac1^-/-^* mutant retinae. Asterisk in (a) indicates unspecific antibody trapping in the lens. Statistical significance was determined using Student’s t-test and depicted as: * p-value < 0.05; ** p-value < 0.01; ns: non significant. Scale bars, 50 μm.

### High levels in *9*-HSA content impact on cell cycle effectors expression in the zebrafish retina

The defects in differentiation and proliferation that we previously observed could be consequences of the deregulation of the expression of key cell cycle regulators. We therefore next examined whether *9*-HSA could affect the expression of genes involved in cell cycle regulation and cellular differentiation events. *C-myc* is a proto-oncogene involved in the regulation of cell proliferation and differentiation (Schreiber-Agus *et al*, 1993). In 2 dpf DMSO control retinae *c-myc* mRNA was detectable in the outermost region of the forming CMZ where RSCs and RPCs are located (Fig. 5a, 5c) (Wan *et al*, 2016). In contrast, in *9*-HSA-treated retinae *c-myc* transcripts were found both in the CMZ and in the central retina (Fig. 5a - a’). We also assessed the expression of two important cell cycle regulator genes: *cyclinD1* (also known as *ccnd1*) and *p27* (Yarden *et al*, 1995; Masai *et al*, 2005). In DMSO treated retinae, *p27* was found expressed in patches all over the neuroepithelium at 2 dpf (Fig. 5b). *p27* expression was found drastically down-regulated in *9*-HSA -treated retinae (Fig. 5b’). Unlike *p27, cyclinD1* expression was strongly up regulated in *9*-HSA -treated retinae with respect to control (Fig. 5c – c’). Indeed in *9*-HSA -treated retinae, *cyclinD1* transcripts were detectable in the entire retina, while in control embryos *cyclinD1* expression was confined to the forming CMZ. Therefore, *9*-HSA can affect the expression of crucial cell cycle regulators and effectors. High levels of *9*-HSA in RPCs causes a deregulation in the expression of these cell cycle regulators, which correlates with the increase in proliferation and differentiation defects observed in these retinae.

**Figure 5.**
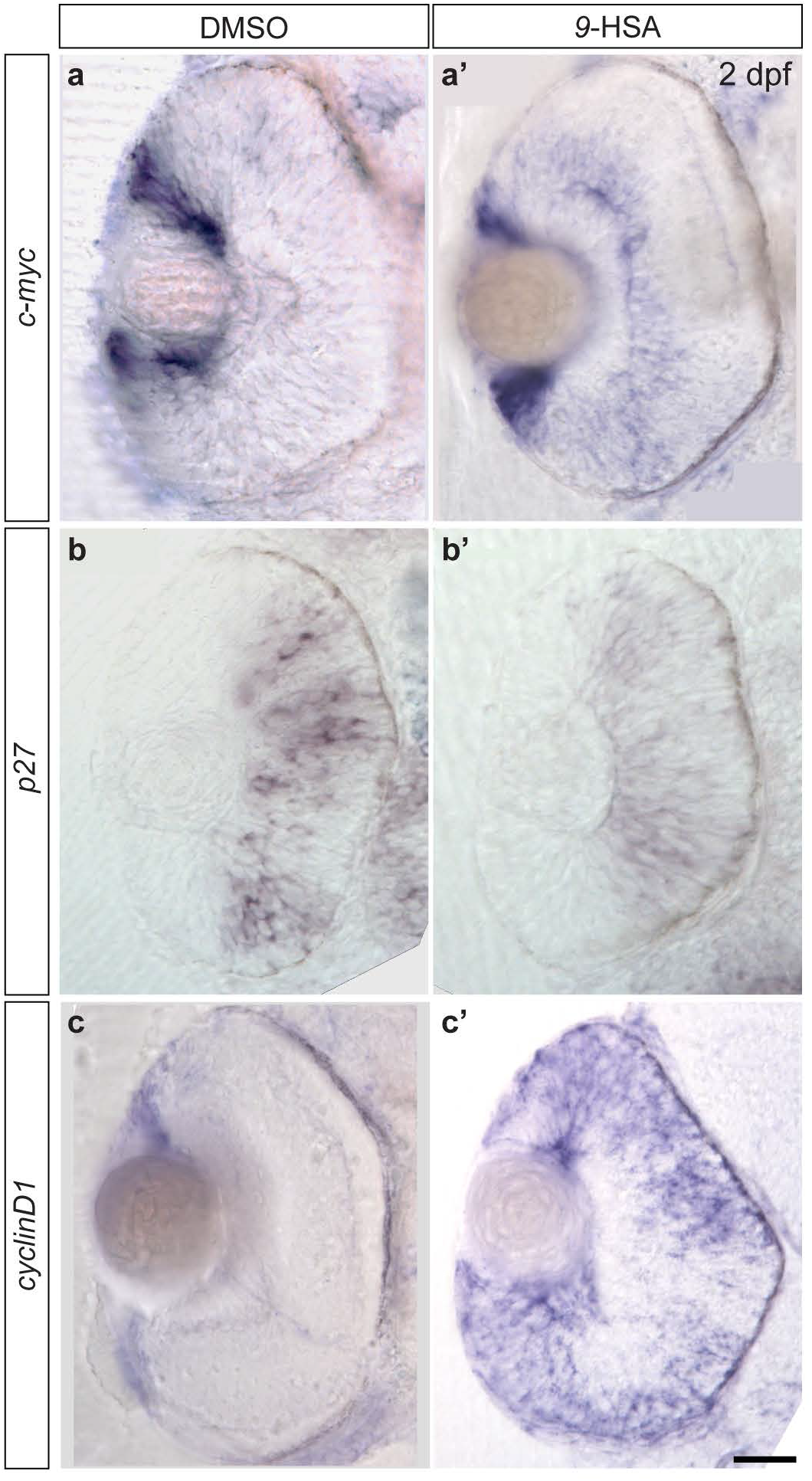
*9*-HSA exogenous administration in retinal progenitor cells affects the expression of cell cycle regulators. Expression analysis of the three cell cycle regulators *c-myc, p27* and *cyclinD1* at 2 days post-fertilization (dpf) in DMSO-treated (respectively a, b, c) and *9*-HSA –treated retinae (respectively a’, b’, c’). (a – a’) Comparative *in situ* hybridization of *c-myc* revealed an increase in its transcription in the central part of *9*-HSA –treated retinae in comparison to DMSO-treated control retinae. (b – b’) Expression analysis of *p27* revealed a complete loss of *p27* expression in *9*-HSA -treated retinae in comparison to DMSO-treated control retinae. (c – c’) Similarly to *c-myc*, comparative analysis of *cyclinD1* expression in DMSO-treated and *9*-HSA –treated retinae showed an increase in the expression of *cyclinD1* in the retinae embryos injected with *9*-HSA in comparison to DMSO control retinae. Scale bar, 50 μm.

### *9*-HSA inhibits HDAC1 activity in vivo in a tissue-specific manner

We previously demonstrated that tumor growth inhibitory effect of *9*-HSA on human colon carcinoma cells (HT29) *in vitro* was mediated through a direct fatty acid interaction, leading to a decrease of the enzymatic activity of the histone deacetylase 1 (HDAC1) (Calonghi *et al*, 2005; Parolin *et al*, 2012). In zebrafish HDAC1 was shown to be required for aspects of the eye and central nervous system development, and for the establishment of neural crest cell populations (Cunliffe, 2004; Cunliffe & Casaccia-Bonnefil, 2006; Harrison *et al*, 2011; Ignatius *et al*, 2013; Stadler *et al*, 2005; Yamaguchi, 2005). In the retina, *hdac1* mutation (known as *add* or *ascending and descending*) results in the failure of RPCs in exiting the cell cycle, correlating with an increase in cell proliferation, cell cycle deregulation, lack of differentiation and therefore absence of proper tissue lamination (Yamaguchi *et al*, 2005). Consistent with these results, we detected defect in retina development fully in line with the effect of *9*-HSA injection (comparing Fig. 3, 4 and 5 with Supplementary Fig. 3, 4 and 5 respectively). These data are strongly suggesting that the effects of *9*-HSA injection are mediated by HDAC1 inhibition in our *in vivo* system. We therefore assessed whether *in vivo*, *9*-HSA can impact on HDAC1 activity. By immunohistochemistry, the histone H4 acetylation (acH4) levels in *hdac1^-/-^* mutant, wild type siblings, *9*-HSA –injected and control (DMSO-injected) zebrafish embryos were evaluated (Fig. 6a – h). Hyper-acetylated H4 levels were strikingly increased at 2 and 3 dpf in *9*-HSA -treated and *hdac1^-/-^* mutant retinae, in comparison to the respective controls (Fig. 6a - h). To further demonstrate that *9*-HSA induces an increase of histone H4 acetylation, a western blot was performed on 3 dpf whole embryo lysates (Fig. 6i). The western blot analysis was also performed to assess the effect of *9*-HSA on the acetylation of lysine 9 of histone H3 (acH3K9, Fig. 6i). A significant increase in the level of histone H4 acetylation could be detected, and the same was observed for the histone H3 acetylation mark (acH3K9) in *9*-HSA -treated and *hdac1^-/-^* mutant embryos with respect to controls. Western blot quantification (n = 3) of H4 and H3 hyper-acetylation marks revealed an increase of 2.7 fold for the histone H4 in *9*-HSA-injected embryos over DMSO-injected embryos (SD ± 1.3) similarly to *hdac1^-/^-* where an increase of 2.8 fold in the acH4 mark was measured (ratio over wild type sibling, SD ± 1.2). Concerning acH3K9 acetylation mark, we measured an increase of 2 fold in *9*-HSA -treated embryos over DMSO-injected embryos (SD ± 0.96) and an increase of 2.3 fold in *hdac1* mutants over wild type sibling (SD ± 0.4) (Fig. 6j). These results therefore indicate that *9*-HSA *in vivo* can act as an inhibitor of HDAC1 and can as such impact on the proliferative capacity of a cell depending on its amounts.

**Figure 6.**
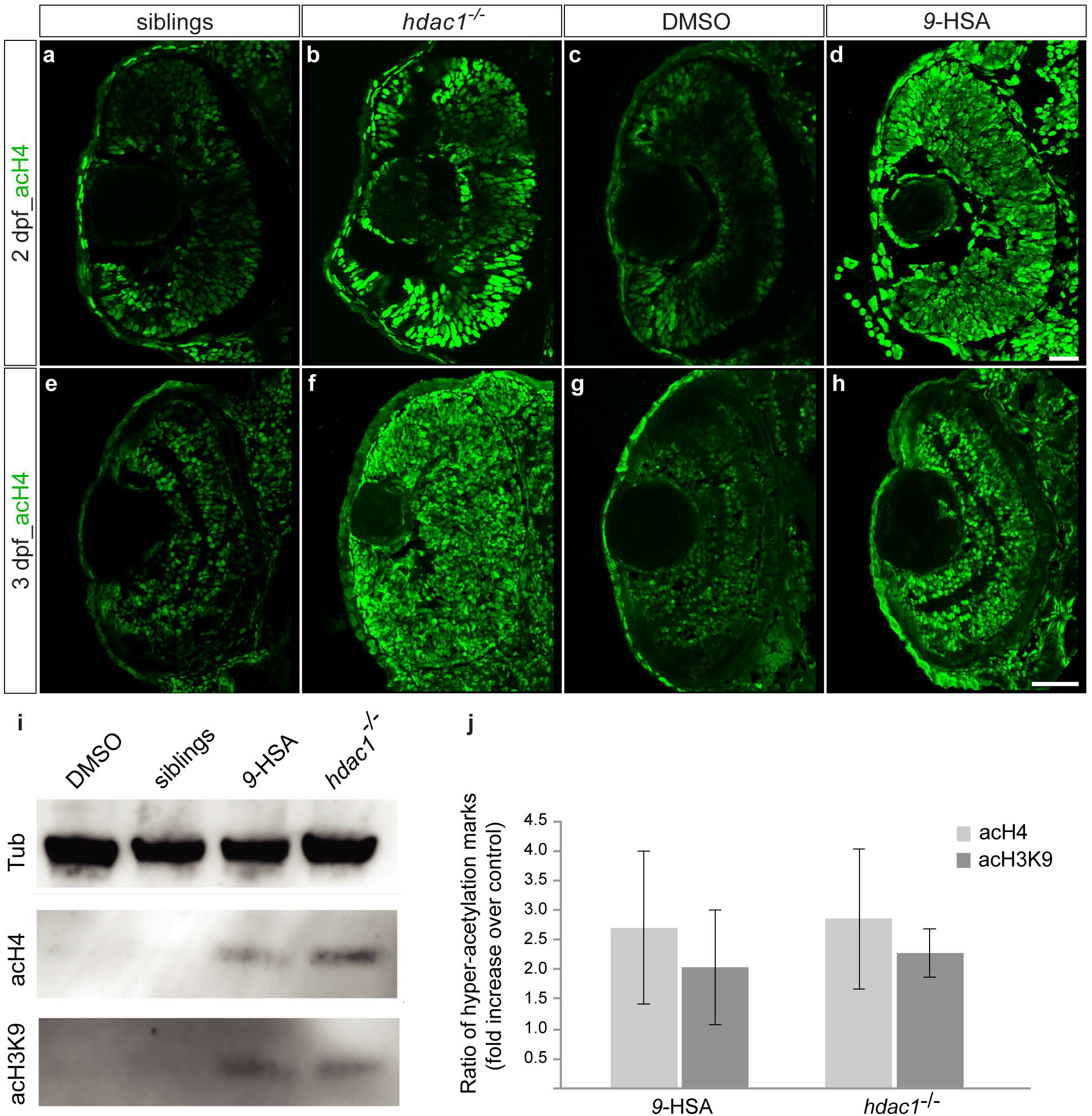
*9*-HSA inhibits HDAC1 enzymatic activity ***in vivo*** in the retina. (a – h) *9*-HSA –treated and *hdac1^-/-^* mutant retinae showed an increase in histone H4 hyper-acetylation marks in comparison to wild type (siblings) and DMSO –treated retinae at 2 and 3 days post-fertilization (dpf). Immunohistochemistry of anti-hyper-acetylated histone H4 (acH4, green) on 2 and 3 dpf frontal retinal cryosections. (i) Western blot analysis of histone H4 and histone H3 hyper-acetylation marks (acH4 and acH3K9 respectively) from 3 dpf DMSO –treated, wild type (siblings), *9*-HSA treated and *hdac1^-/-^* whole embryo lysates showed an increase in both marks in *9*-HSA treated and *hdac1^-/-^* whole embryo lysates. (j) Western blot quantification of hyper-acetylation of the histone 4 (acH4) and histone 3 (acH3K9) marks. Data are presented as ratios of *9*-HSA treated over DMSO –treated embryos and *hdac1^-/-^* mutant over wild type (siblings). Values are means ± standard error of three independent samples. Scale bar (a – d), 20 μm; scale bar (e – h), 50 μm.

In order to assess the specificity of *9*-HSA in promoting RPCs proliferation, we evaluated its effect on the development of the hindbrain where HDAC1 enzymatic activity was shown to play an opposite role with respect to its retinal function (Cunliffe, 2004; Stadler *et al*, 2005; Yamaguchi, 2005). Indeed in this tissue, HDAC1 was shown to be required for the responsiveness of precursors cells to Hedgehog signaling pathway signals by repressing Notch signaling pathway effectors and its loss-of-function leads to a reduction of cell proliferation at 1 dpf (Cunliffe, 2004). In line with these results, we observed a significant reduction in cell proliferation in the hindbrain of 1 dpf *9*-HSA -treated in comparison to DMSO control embryos (n = 7 hindbrains for DMSO and n = 9 for *9*-HSA treated hindbrain. Average number of pH3-positive cells in DMSO control hindbrain = 246, SD ± 26 and average number of pH3-positive cells in DMSO control hindbrain = 203, SD ± 25; p-value: 0.005142, Supplementary Fig. 6A - C). Unlike in the retina where *hdac1* loss-of-function and *9*-HSA exogenous administration both lead to an increase in proliferation (Fig. 4d), in the hindbrain, a decrease in the number of proliferating cells can be observed after *9*-HSA treatment. Therefore the observed effects of *9*-HSA *in vivo* on proliferation are tissue-specific and its regulation of progenitor/precursor cell fates are fully consistent with the inhibition of HDAC1 activity and cannot be explained by a general developmental delay. By increasing its endogenous levels, we reveal that *9*-HSA *via* HDAC1 inhibition can regulate the epigenetic state of RPCs, placing HDAC1 as a downstream target of the lipid peroxidation process.

### H_2_O_2_ degradation by Catalase is sufficient to induce RPCs differentiation

Together our results highlight the crucial role for H_2_O_2_ and lipid peroxidation downstream products like *9*-HSA in regulating progenitor cell fate. In order to test our model, we locally perturbed H_2_O_2_ levels by clonally over-expressing *catalase* under the control of *rx2* promoter and evaluated the effects on RPC (Fig. 7a). To show the activity of our construct on endogenous H_2_O_2_ levels in the developing neuroepithelium, we injected *UAS:catalase-E2A-tagRFP* in the double transgenic *Tg(ubi:HyPer),Tg(rx2:gal4).* We imaged the injected retinae at 24 hpf when H_2_O_2_ levels are high and homogeneous throughout the retinal epithelium (Fig. 1a). We observed that in the RFP-positive clones expressing ectopic *catalase*, H_2_O_2_ levels were decreased with respect to their neighbor cells, showing that H_2_O_2_ levels can be modulated by our construct in a temporal and local manner (Figure 6b). We then decided to evaluate the effect of the decrease in the levels of H_2_O_2_ in those progenitors. To do so, we fixed *UAS:catalase-E2A-tagRFP* -injected and *UAS:tagRFP* control injected embryos at 48 hpf and counterstained them with the photoreceptors marker Zpr1. Photoreceptor differentiation starts at 48 hpf in a small patch of cells in the ventro-central part of the retina and by 3 dpf, Zpr1-positive cells are found throughout the photoreceptor cell layer (Liu *et al*, 2007). We focused our analysis looking at the RFP-positive clones and derived differentiated cells located at the dorsal periphery of the retinal tissue where cells do not yet express Zpr1 as could be observed in *UAS:tagRFP* control injected embryos (Fig. 7c and d). Surprisingly, the early expression of *catalase* induced in rx2-expressing clones was sufficient to drive the differentiation of Zpr1-positive cells in all clones examined (Fig. 7c and d). Together these results show that, upstream of lipid peroxidation events, the levels of H_2_O_2_ and their degradation by scavengers like Catalase are critical to trigger the switch from proliferation to differentiation of progenitors cells during retinal development.

**Figure 7.**
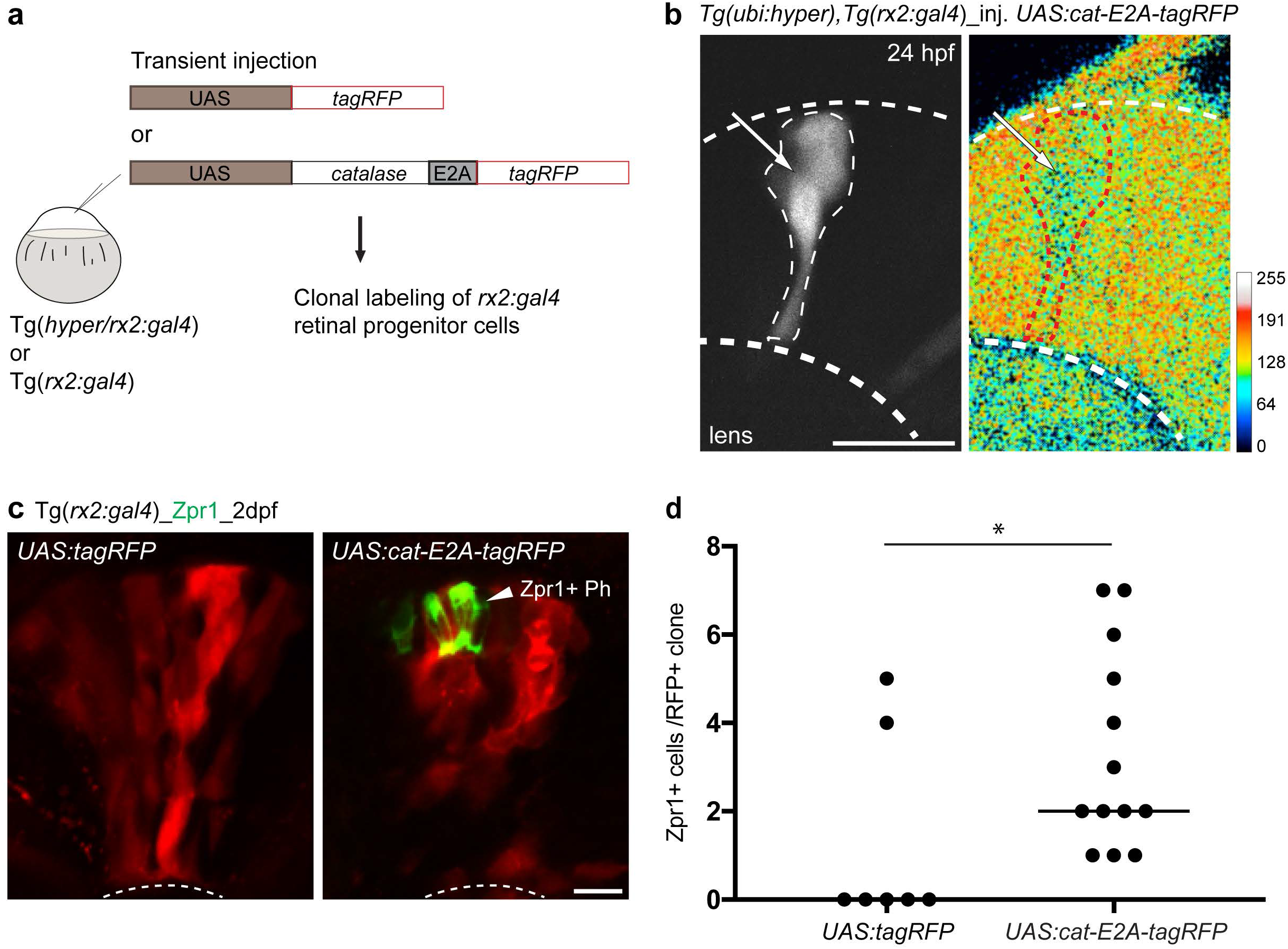
Ectopic expression of ***catalase*** in RPCs is sufficient to trigger premature differentiation. (a) Cloning strategy of Catalase transient and mosaic overexpression in *rx2:gal4*-expressing RPCs with or without *Tg(ubi:HyPer)* transgenic background. (b) Early *catalase* overexpression in *rx2-* expressing RPCs (white arrow) can decrease the levels of H_2_O_2_ in these progenitors at 24 hours post-fertilization (hpf) as revealed by the *HyPer* transgene reporting the endogenous levels of H_2_O_2_ of *UAS:catalase-E2A-tagRFP (UAS:cat-E2A-tagRFP*) injected double transgenic *Tg(ubi:HyPer),Tg(rx2:gal4)* embryos (white arrow and red dash line). (c) Early *catalase* overexpression by injection of *UAS:catalase-E2A-tagRFP* in *Tg(rx2:gal4*) RPCs led to the premature differentiation of their deriving cells as revealed by the late photoreceptor marker Zpr1 (white arrowhead, Zpr1+ Ph) in contrast to control *UAS:tagRFP*-injected RPCs. (d) Quantification of Zpr1-positive cells in *UAS:tagRFP* control clones and *UAS:cat-E2A-tagRFP* expressing clones revealed a significant increase in the number of Zpr1-positive cells in clones that expressed *catalase* (n = 7 retinae for *UAS:tagRFP* control and n = 13 retinae for *UAS:cat-E2A-tagRFP, p-value* = 0.02, Mann-Whitney U test). Medians for both conditions are shown (0 for *UAS:tagRFP* control and 2 at *UAS:cat-E2A-tagRFP).* Scale bars, 10 mm.

## Discussion

Our understanding of the role of ROS in general and H_2_O_2_ in particular has evolved in the last years. It is now evident that far from being just detoxification by-products of harmful processes these metabolites are involved in cell signaling during embryonic development, cell growth and stem cell fate regulation and regeneration (Bigarella *et al*, 2014; Zhou *et al*, 2014; Gauron *et al*, 2016; Le Belle *et al*, 2011; Hervera *et al*, 2018; Meda *et al*, 2017; Gauron *et al*, 2013; Sies, 2017; Sies *et al*, 2017). More recently, the role of ROS started to be widely investigated in stem cell field, highlighting the inverse correlation between their amount and stem cells quiescence. The shift in the metabolic balance of these cells between glycolysis and oxidative phosphorylation is linked to the levels of ROS generated and degraded in their niche and the state in which the cells are, quiescent, pluripotent or differentiating progenitor cells (Shyh-Chang *et al*, 2013). Therefore the regulation of this shift “*plays a pivotal role in dictating whether a cell proliferates, differentiates or remains quiescent*” (Shyh-Chang *et al*, 2013). Zhou et al. demonstrated a role for ROS in the generation of pluripotent cells acting on the nuclear reprogramming of somatic cells (Zhou *et al*, 2016). They indeed showed that the onset of reprogramming of somatic cells is tightly associated with an increase in ROS production, which need to be at optimal levels for the process to efficiently occur. They also observed that this initial increase in the production of ROS in these cells is later decreased and maintained at lower levels by the up-regulation of ROS scavenger enzymes and antioxidants such as Catalase or Superoxide Dismutase, preventing these cells from cellular and genomic damages triggered by excessive ROS exposure. Therefore the levels of ROS and their regulation are essential to ensure their physiological function. We have previously shown that *catalase* is expressed specifically in the developing central nervous system of zebrafish, suggesting that a similar mechanism may be in place during normal neurogenesis (Gauron *et al*, 2016). Interestingly, using *Xenopus* retinal explants Agathocleous et al. revealed that glycolysis is an essential process in the CMZ possibly relevant for the quiescent, stem and/or progenitor cells that are all present in the niche (Agathocleous *et al*, 2012). This study in particular suggests that the regulation of glycolysis and oxidative phosphorylation is critical for normal retinal development.

Here we were able to reveal the dynamics of H_2_O_2_ levels in the developing zebrafish retina, a tissue with clear spatial compartmentalization separating quiescent, proliferative, cycling progenitor and differentiated cells. We could draw for the first time a clear correlation between the proliferative state of the neuroepithelium and levels of H_2_O_2_. From these observations, we decided to get further insight on the regulation of this dynamics and assess the spatio-temporal expression of *catalase* in this tissue. The expression of *catalase* and the pro-neural gene *ath5* coincide at 2 dpf across the central retina and later, at 3 dpf, in the CMZ where they are both maintained (Kay *et al*, 2005). Analyzing *catalase* expression in *ath5*÷ mutants, we demonstrate that it is independent from Ath5 function itself (Kay *et al*, 2001; Masai *et al*, 2000). This suggests that Catalase regulation and function act in parallel to the pro-neuronal activity of Ath5 in committed RPCs. In contrast, the presence of the end product of the lipid peroxidation 4-HNE in the domain of Rx2 in the CMZ where RSC and multipotent RPCs are located indicates that lipid peroxidation by ROS is active in these cells and can be involved in their proliferative capacity. In this context, the down-regulation of ROS levels by Catalase can act as a trigger to start the cascade leading to neural differentiation.

To unravel the role of ROS scavenging and lipid peroxidation in RPCs, we investigated the *in vivo* function of *9*-HSA, a known downstream product of lipid peroxidation that we previously demonstrated as a regulator of cellular growth in different human tumor cell lines (Bertucci *et al*, 2002; Calonghi *et al*, 2007; Cavalli *et al*, 1991; Gesmundo *et al*, 1994). In these studies, we revealed by biochemical and structural modeling studies that, *in vitro*, *9*-HSA inhibits histone deacetylase 1 (HDAC1) activity by directly binding to its catalytic site (Calonghi *et al*, 2005; Parolin *et al*, 2012). Both *9*-HSA enantiomeric forms *(R*) and (S)-*9*-HSA have the ability to inhibit HDAC1 activity, with (*R*) being more active, through a direct ligand-enzyme interaction, thereby blocking substrate access (Calonghi *et al*, 2005). The acetylation and deacetylation of core histones of chromatin are essential histone modifications required for many biological processes such as cell proliferation, differentiation and gene silencing (Allis & Jenuwein, 2016). While *9*-HSA role *in vivo* had not been characterized so far, HDAC1 was shown to play a critical function in regulating the balance between proliferation and differentiation, and this has been reported in many different neuronal tissue in zebrafish (Yamaguchi *et al*, 2005; Cunliffe, 2004; He *et al*, 2016). However, the upstream events leading to HDAC1 activity modulation remained unknown. Here by mass spectrometry we were able to detect *9*-HSA in the zebrafish embryo, indicating that the molecule is endogenously produced during zebrafish development. By exogenous administration, we were able to evaluate the effect of *9*-HSA overexposure on RPCs. In contrast to its role in cancer cell lines, we show that high amounts of *9*-HSA in the retina are able to alter the balance between cell proliferation and differentiation and reveal that, like *in vitro*, *9*-HSA acts as an endogenous inhibitor of HDAC1 *in vivo*, therefore regulating the epigenetic state of RPCs. In the zebrafish retina HDAC1 was shown to be required to promote differentiation of RPCs through the inhibition of Notch and Wnt signaling pathways (Yamaguchi, 2005). In zebrafish and amphibian retinae Wnt signaling has been found to promote proliferation while Notch prevents differentiation (Borday *et al*, 2012; Denayer *et al*, 2008; Del Bene *et al*, 2008; Schmidt *et al*, 2013; Yamaguchi, 2005). Interestingly, in contrast to its proliferative effect in the retina, *9*-HSA has an opposite effect in the hindbrain where we observed a decrease in the mitotic index. This is in agreement with the reported function of HDAC1 in this tissue (Cunliffe, 2004). Together our results indicate a context-dependent modality of action of *9*-HSA, which *via* HDAC1 inhibition, can differentially influence stem and progenitor cell fate in a tissue-specific manner, and demonstrate that its effect on retina proliferation cannot be explained by simple embryonic developmental delay. Overall our data identify *9*-HSA as a possible signaling molecule acting downstream H_2_O_2_ and contributing to the fine regulation of the balance between proliferation and differentiation in neuronal progenitors. In this context, we propose *9*-HSA to work in parallel with other well-characterized H_2_O_2_ related signaling event acting on cysteine redox state on a number of proteins (Go *et al*, 2015; Yang *et al*, 2016).

To test our model and gain further insight in the upstream regulation of *9*-HSA production, we next decided to locally interfere with H_2_O_2_ levels in RPCs. To do so, we clonally introduced the expression of Catalase in RPCs at a stage when the epithelium is homogeneously enriched in H_2_O_2_. We demonstrated the scavenging activity of Catalase in these clones and could show that decreasing the endogenous levels of H_2_O_2_ at this stage was sufficient as a signal for RPCs to initiate their differentiation. These observations demonstrate the importance of the fine-tuning of the physiological redox signaling, here through H_2_O_2_ and lipid peroxidation regulation in RPCs.

Taken together our work reveal the critical and physiological function of the redox signaling and lipid metabolism products in the retina and the importance of their fine-regulation. These products, like *9*-HSA, can act as endogenous intermediate modulators allowing RPC to initiate differentiation during retinal development and later in the maintenance of tissue homeostasis in the adult retina. In addition to this new role in the retina, other work on derivatives of the lipid such as fatty acid esters and in particular palmitic acid esters of *9*-HSA (*9*-PAHSA) demonstrated their in anti-diabetic and antiinflammatory properties, extending its role to another range of biological activities (Yore *et al*, 2014; Lee *et al*, 2016; Nelson *et al*, 2017). Our study provides insights in the mechanism of action of *9*-HSA *in vivo via* its inhibitory activity on HDAC1 and proposes a novel mechanism linking H_2_O_2_ homeostasis and neuronal differentiation *via* the modulation of lipid peroxidation that leads to the expression of key neuronal genes required for the switch from proliferation to differentiation of RPCs (Fig. 8).

**Figure 8.**
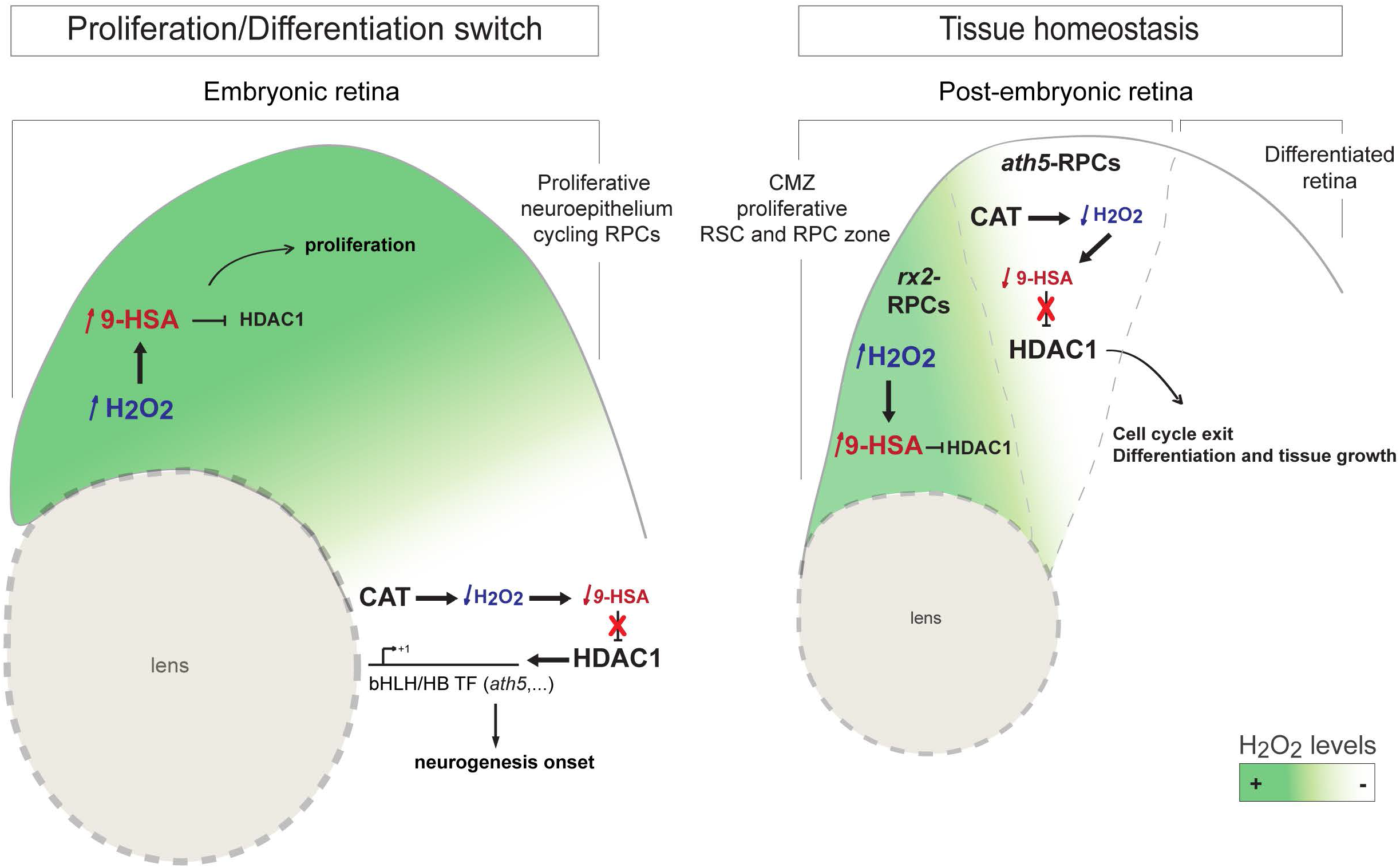
Schematic modeling of the cascade of events linking ROS scavenging/lipid peroxidation to retinogenesis and the maintenance of retinal tissue homeostasis. The embryonic retina is initially comprised of cycling retinal progenitors cells (RPCs), which initiates retinal differentiation upon *ath5* expression in the central part of the retina. Λth5-expressing RPCs also express *catalase* and are therefore able to respond to high ROS and lipid peroxidation product generation, including *9*-HSA. Catalase up-regulation inhibiting lipid peroxidation lifts the inhibition on HDAC1, which promotes differentiation in the retina. In the post-embryonic fish retina, cycling progenitors are located in the ciliary marginal zone (CMZ). The high levels of ROS and lipid peroxidation products in the niche as well as the expression of *catalase* within the domain of Ath5 suggest a conserved mechanism in the regulation of oxidative stress between embryonic and post-embryonic RPCs, which can maintain the homeostasis of the retinal tissue.

## Material and methods

### Zebrafish stocks and ethic statements

All fish are housed in the fish facility of our laboratory, which was built according to the local animal welfare standards. All animal procedures were performed in accordance with French and European Union animal welfare guidelines. *Hdac1\** mutant fish used in this study were raised from embryos kindly supplied by the Masai laboratory.

### Transgenic fish lines, micro-injection and imaging

Imaging and ratiometric analysis of HyPer signal in this line and double transgenic *Tg(ubi:HyPer),Tg(rx2:gal4)* was performed as described in (Gauron *et al*, 2016). *Tg(rx2:gal4*) line was previously described in (Di Donato *et al*, 2016). Except for the imaging of *Tg(ubi:HyPer)* and *Tg(ubi:HyPer),Tg(rx2:gal4)*, Zeiss LSM780 and LSM880 confocal microscopes (Zeiss) were used for high resolution microscopy, employing a 40× water immersion objective. Embryos were embedded in 1% low melting agarose and Z-stacks were acquired every 1- to 2-μm.

### Molecular cloning

The *pUAS:tagRFP* construct used in this study was previously cloned in (Auer & Del Bene, 2014). For the generation of the *pUAS:catalase-E2A-tagRFP* plasmid full cDNA of *catalase* was PCR amplified from total cDNA of 2 dpf zebrafish embryos and the *tagRFP* sequence amplified from the same *pUAS:tagRFP* plasmid. The two fragments, separated by a E2A cassette were cloned into the *pUAS:ubc-pA* plasmid (Horstick *et al*, 2015) with the Gibson Assembly Cloning Kit.

### Synthesis of *9*-HSA

General ^1^H and ^13^C NMR spectra were recorded on a Mercury 400 (or an Inova 600) spectrometer (Varian, Palo Alto, CA, USA) at 400 (or 600) MHz and 100.57 (or 150.82) MHz, respectively. Chemical shifts were measured in δ, and referenced to TMS for ^1^H NMR and to CDCL (77.0 ppm for ^13^C NMR). *J* values are given in Hz. Signal multiplicities were established by DEPT experiments. Mass spectrometry was performed using a VG-7070E spectrometer at an ionisation voltage of 70 eV. IR spectra were recorded using a Perkin-Elmer FT-IR MOD.1600 spectrophotometer. Melting points were determined with a Büchi 535 or a Stuart SMP3 apparatus and are uncorrected. Catalytic hydrogenation was performed using a Parr 3911 shaker type hydrogenator (Parr Instrument Company, Moline, Illinois, USA). Silica gel 230-400 mesh and silica gel coated plates Kieselgel 60 F254 were purchased from Merck (Darmstadt, Germany) and were used for flash-chromatography (FC) and TLC, respectively, the spots being developed with an aqueous solution of (NH4)MoO4 (2.5%), (NH4)4Ce(SO4)4 (1%) in 10% ¾SO4. Preparative TLC was performed by using 20 x 20 cm silica gel coated plates (Merck, Darmstadt, Germany). *Dimorphotheca sinuata* L. seeds were purchased from Galassi Sementi (Gambettola, Forlì-Cesena, Italy). (*R*)-(-)-O-acetyl mandelic acid was from Fluka Chemie GmbH (Buchs, Switzerland). Anhydrous dichloromethane was obtained by distillation over P_2_O_5_. Racemic *9*-HSA was prepared as previously reported (Calonghi *et al*, 2005), as well as 10-HSA (Ebert *et al*, 2012). Pure (*R*)-*9*-HSA was prepared from *Dimorphotheca sinuata* seeds according to the procedure previously described (Parolin *et al*, 2012; Boanini *et al*, 2016) and its optical purity (>99%) was determined by H NMR analysis after derivatization with (*R*)-(-)-O-acetylmandelic acid (Parolin *et al*, 2012). The synthesized (*R*)-*9*-HSA enantiomere, previously shown to be more efficient in inhibiting HDAC1 activity was used for exogenous administration (Parolin *et al*, 2012).

### Micro-injections

*9*-HSA was solubilized in dimethylsulfoxide (DMSO, Sigma Aldrich), to obtain a solution with a final concentration of 33 mM. For the control embryos, pure DMSO was used. To each solution 0.1% of Phenol red solution was added in order to control the amount of solution injected, and an average volume of 1 nL was injected in each embryo. Embryos were injected into the yolk about 10 - 15 minutes after fertilization, between zygote and 4-cells stage.

*UAS:tagRFP* and *UAS:catalase-E2A-tagRFP* plasmids were injected at 1-cell stage into *Tg(rx2:gal4*) and double transgenic *Tgubi:(HyPer),Tg(rx2:gal4)* lines. 50 ng/μL of *tol2* and *tol1* mRNA were co-injected with 20 ng/μL of *UAS:tagRFP* and *UAS:catalase-E2A-tagRFP* plasmids respectively.

### LC-ESI-MS analysis of lipid extract

A total lipid extract was obtained from embryos from day 0 to day 5. For each time point two hundred wild-type embryos were used and processed using the Folch method (Folch *et al*, 1957). Stock standard solution was prepared by dissolving *9*-HSA and 10-HSA at a concentration of 2 mg/mL in methanol. Further dilutions for the calibration curve were prepared daily in methanol:0.05% acetic acid in water (80:20), (v/v). A calibration curve for the quantitative determination of *9*-HSA was built in the range of 0.2222 *μ*g/mL. Samples were solubilized with 500 *μ*l of methanol:0.05% acetic acid in water (80:20), (v/v), sonicated and filtered with a 0.45 *μ*m syringe filter. Chromatographic analysis were performed with a Sunfire C18 (2.1×100 mm, 3.5*μ*m; Waters) using the following mobile phases: water with 0.05% acetic acid (A) and methanol with 0.05% acetic acid (B). The gradient elution started with 70% of solvent B, reached 90% of solvent B in 15 minutes, remained at 90% for 5 minutes and then returned to 70% of solvent B in ten minutes. The column temperature was set at 30°C. The injection volume was 2 *μ*L and the flow rate was 0.2 mL/min. LC-MS/MS analysis was carried out using a Series200 modular HPLC (PerkinElmer) interfaced with an API 4000-QTrap triple quadrupole (AB-Sciex Framingham, MA, USA) with an electrospray ionization (ESI) source. MS parameters were optimized by infusing standard solutions at a concentration of 2.2 *μ*g/ml into the TurboIonSpray1 source operating in negative ion mode by a syringe infusion pump.

The optimized parameters of the ion source were: 600°C probe temperature, 30 psi curtain gas, 40 psi nebulizing gas, 40 psi heating gas, −4000 kV ion spray voltage. For quantitative analysis mass chromatograms were acquired in multiple reaction monitoring (MRM) in double. Parent ion / daughter ion mass-to-charge quantitative and qualitative transition for *9*-HSA were 299.1/253.2 and 299.1/155.0m/z, declustering potential, entrance potential, collision energy and cell exit potential for quantitative and qualitative transitions were: 70V, −10V, −35eV, −10V and −70V, −10V, −35eV, −8V, respectively. Data processing and quantitation were performed by Analyst software (AB-Sciex). Calibration was done by linear regression with a 1/x weighting; and analyte concentration in each sample was back calculated by the interpolation on the regression curve.

### Immunostainings

Immunohistochemistry experiments were performed as previously described (Masai *et al*, 2000). The antibodies used in this study and their dilutions were the following: anti-hyperacetylated histone H4 (1:500, Penta 06-946, Millipore), anti-phospho-histone H3 (1:500, 06-570, Millipore), anti-PCNA antibody clone PC10 (1:500, AB477413, Sigma Aldrich), anti-Zn-5 and Zpr-1 (1:500, Zebrafish International Resource Center), anti-parvalbumin (1:500, MAB1572, Millipore), anti-glutamine synthetase (1:500, MAB302, Millipore) and anti-4-hydroxynonenal (#HNE13-M, Interchim). Secondary antibodies used in this study were: Alexa546 fluorophore-conjugated secondary antibody (1:500, Invitrogen Molecular Probes), Alexa488 fluorophore-conjugated secondary antibody (1:500, Invitrogen Molecular Probes), DAPI for nuclear labeling (1:500, Sigma).

### TUNEL assay

Apoptosis was detected in whole zebrafish embryos at 3 dpf by TUNEL assay, using Apoptag Peroxidase In Situ Apoptosis Detection Kit (Millipore). The embryos were fixed in 4% PFA for 2 hours at room temperature and then stored in methanol at −20°C and processed according to the standard protocol. After the staining, the embryos were embedded in gelatin/albumin with 4% glutaraldehyde, and 20 μm retinal sections were generated using a VT1000 S vibrating blade microtome (Leica). The sections were then analyzed on a Leica Upright Widefield epifluorescence microscope.

### BrdU incorporation

Embryos 2 dpf were dechorionated and anesthetized with 1mM MS-222 (TMS, tricaine methanesulfonate solution, Sigma), followed by BrdU (Sigma) injection into the yolk. The injection solution was composed by a BrdU at the final concentration of 50 mM, Phenol Red at the final concentration of 0.5% in ddH2O. 12 hours after the BrdU injection, the embryos were washed, fixed in 4% PFA and cryo-sectioned (Leica) with a thickness of 12 μm. Sections were then treated with a denaturing 2N HCl solution for 20 minutes at 37°C followed by five PBS washing steps. After a blocking of 2 hours in 5% normal goat serum in PBTD (PBT + DMSO), the sections were incubated with unconjugated anti-bromodeoxyuridine antibody (1:1000, PRB1-U, Phoenix) in block solution overnight at 4°C. The slides were then washed with PBTD and incubated with Alexa546 fluorophore-conjugated secondary antibody (1:500, Invitrogen Molecular Probes) diluted in blocking solution for 1.5 hours at room temperature. The slides were finally stained with the nuclear marker DAPI (1:500, Sigma) for 5 minutes at room temperature.

### Western blot analysis and quantification

Yolk-extirpated embryos from different time points were homogenized using an extraction buffer (50 mM Tris-HCl, pH 6.8; 2% SDS; 10% glycerol; 12% 2-mercaptoethanol) to extract proteins. The extracted proteins were subjected to SDS-PAGE and blotted onto a PVDF membrane (BioRad). After blocking with 5% skim milk and 0.1% Triton X in TBS, membrane filters were incubated with an anti-hyperacetylated histone H4 (Penta, 06-946, Millipore) or anti-acetylH3K9 antibody (Abcam, Ab4441). Immuno-signals were visualized with an alkaline-phosphatase (AP)-conjugated anti-rabbit IgG antibody (1:5000, Promega). Western blot revelation was performed using a cheminoluminescence digital imaging system (ImageQuant Las-4000 Mini, GE Healthcare Life Sciences) and quantified using ImageJ software.

### Whole-mount single and double fluorescent in situ hybridization

Digoxigenin-labeled riboprobes were prepared as recommended by the manufacturer instructions (Roche). Whole-mount *in situ* hybridization was performed using standard procedures (Oxtoby & Jowett, 1993). Further details of the probes used are available upon request. For double fluorescent *in situ* hybridization, standard digoxigenin and fluorescein-conjugated riboprobes were revealed using Tyramide Signal Amplification (TSA) kits (PerkinElmer). *Ath5* riboprobe was incubated for 40 minutes and *catalase* riboprobe for 60 minutes, after which embryos were kept in the dark and washed with a TNT solution (0.1 m Tris, pH 7.5, 0.15 m NaCl, 0.1% Tween 20). Following these washes, embryos were incubated for 20 minutes in 1% H_2_O_2_ in TNT, washed several times and blocked with TNB [2% DIG Block (Roche) in TNT] for 1 hour. These steps were followed by an incubation of the embryos in anti-digoxigenin-POD (peroxidase) Fab fragments (Roche, 1:50 in TNB). The riboprobe signals were finally detected using fluorescein-tyramide (FITC) and cyanine3-tyramide (Cy3). The stained embryos were lastly incubated in DAPI in TNT overnight at 4°C and washed with TNT.

### Cryosections and vibratome sections

Cryosections were performed as follows: embryos were fixed in 4% paraformaldehyde in PBS (pH 7.4) for 2 hours at room temperature and cryoprotected overnight in a 30% sucrose/0.02% sodium azide/PBS solution. Embryos were then transferred to plastic molds and embedded in OCT. The blocks were placed on dry ice before sectioning. The sections were cut with a thickness of 12-14 μm and mounted on Fisherbrand Superfrost plus slides (No. 12-550-15). For vibratome sections, whole-mount embryos were washed twice in 1x *PBS*/0.1% *Tween-20* (PBS-Tw) solution. The samples were embedded in gelatin/ albumin with 4% of glutaraldehyde and sectioned (20 μm) on a VT1000 S vibrating blade microtome (Leica). Sections were mounted in Fluoromount Aqueous Monting Medium (Sigma).

### Quantification and image processing

Quantifications within the central retina were generated as described in (Yamaguchi, 2005). A statistical t-test was performed for pH3 quantifications. For Zpr1 cells, a Mann-Whitney U test was used. All images were analyzed using ImageJ, Volocity, Adobe Photoshop and Adobe Illustrator softwares.

## Acknowledgments

We are grateful to Hichiro Masai for sharing reagents and mutant lines. We thank all members of the Del Bene lab for fruitful discussions and inputs. We also thank Marion Thauvin from the Vriz lab for her help in imaging and ratiometric analysis of the HyPer signal of the *Tg(ubi:HyPer),Tg(rx2:gal4)* embryos. We thank the Developmental Biology Curie imaging facility (PICT-IBiSA@BDD, Paris, France, UMR3215/U934) member of the France-BioImaging national research infrastructure for their help and advice with confocal microscopy. The Del Bene laboratory ‘‘Neural Circuits Development’’ is part of the Laboratoire d’Excellence (LABEX) entitled DEEP (ANR-11-LABX-0044), and of the École des Neurosciences de Paris Ile-de-France network. S.A. was supported by a postdoctoral fellowship from the foundation Berthe Fouassier (Fondation de France #201500060246 and #201600069985). This work has been supported by an ATIP/AVENIR program starting grant (F.D.B.), ERC-StG #311159-Zebratectum (F.D.B.), CNRS, INSERM, Institut Curie (F.D.B) and core funding. F.N. was supported by funds from the LABEX (ANR - 11-LABX-0044), a visitor fellowship of the Fondation Pierre-Gilles de Gennes pour la recherche, travelling fellowships of European Network on Fish Biomedical Models, Consorzio Interuniversitario Biotecnologie, Marco Polo project (University of Bologna). C.P. was supported by a STSM Grant from the COST Action BM0804. N.C and C.B were supported by the RFO FUNDS (University of Bologna). This work received support under the programme “Investissements d’Avenir” launched by the French Government and implemented by the ANR with the following references: ANR-10-LABX-54 MEMO LIFE – ANR-11-IDEX-0001-02 PSL* ResearchUniversity.

## Author contributions

S.A. and F.N. designed, performed experiments and analyzed the data with the technical help of K.D. C.G. built the *Tg(ubi:HyPer)* transgenic fish and took HyPer images. C.P. performed pH3 immunostaining on the hindbrain as well as quantification. C.B. performed *9*-HSA syntheses and J.F. the endogenous detection of *9*-HSA. S.A., F.N. and F.D.B. wrote the manuscript with the critical input from all other authors. SV, F.D.B and N.C. conceived and supervised the study.

