## supplemental files for "Redox signaling *via* lipid peroxidation regulates retinal progenitor cell differentiation"

### Supplementary information

**Supplementary Figure 1. *Catalase* expression is independent from Ath5 function in the retina.** Comparative expression analysis of *catalase* expression in wild type (**A**) and *ath5*$^{-/-}$ mutant retinae (**B**) reveals that *catalase* expression is maintained at 3 dpf in the ciliary marginal zone (asterisks, CMZ). Scale bar (**A – B**), 100 µm.

**Supplementary Figure 2. 9-HSA exogenous administration does not trigger apoptosis in the retina at 3 days post-fertilization.** DMSO –treated control (**A**) and *9*-HSA –treated retinae (**B**) show no difference in the number of apoptotic cells at 3 days post-fertilization (dpf) as revealed by TUNEL assay. Scale bar, 20 µm.

**Supplementary Figure 3. *Hdac1*$^{-/-}$ mutant retinae display differentiation defects for all retinal cell types.** (**A – D'**) *Hdac1* loss-of-function results in a lack of differentiation in all cell types analyzed. (**A – A'**) Immunohistochemistry of anti-Zn5 (green) labeling retinal ganglion cells (RGC) on 2 days post-fertilization (dpf) frontal retinal cryosections from wild type (sibling, **A**) and *hdac1*$^{-/-}$ mutant (**A'**) embryos. No RGC labeling could be observed in the central part of the retina at 2 dpf in *9*-HSA -treated and *hdac1*$^{-/-}$ mutant retinae in comparison to wild type and DMSO-treated control retinae. (**B – B'**) Immunohistochemistry of anti-Parvalbumin (green) labeling amacrine cells and displaced amacrine cells (AC) on 3 dpf frontal retinal cryosections from wild type (sibling, **B**) and *hdac1*$^{-/-}$ mutant (**B'**) embryos. Similarly to RGCs, no amacrine cells or displaced amacrine cells could be observed in *hdac1*$^{-/-}$ mutant retinae in comparison to wild type retinae. (**C – C'**) Immunohistochemistry of anti-GS (green) labeling Müller glia cells (MG) on 3 dpf frontal retinal cryosections from wild type (sibling, **C**) and *hdac1*$^{-/-}$ mutant (**C'**) embryos shows no MG differentiation in *hdac1*$^{-/-}$ mutant retinae in comparison to the control retinae. (**D – D'**) Immunohistochemistry of anti-Zpr1 (green) labeling photoreceptor cells (Ph) on 3 dpf frontal retinal cryosections of wild type (sibling, **D**) and *hdac1*$^{-/-}$ mutant (**D'**) embryos. No Zpr1 labeling could be observed in *hdac1*$^{-/-}$ mutant retinae, while in wild type retinae Zpr1-positive Ph could be detected within the outer nuclear layer of the 3 dpf retinal tissue. Scale bar, 50 µm.

**Supplementary Figure 4. *Hdac1*$^{-/-}$ retinae show an increase in proliferation.** (**A – A'**) Immunohistochemistry of anti-PCNA (red) labeling cells in S-phase of the cell cycle on 3 dpf frontal retinal cryosections. *Hdac1*$^{-/-}$ retinae show an expanded PCNA labeling in the ciliary marginal zone (CMZ, bracket) and in the central retina in comparison to wild type (siblings) retinae at 3 dpf. (**B – B'**) Immunohistochemistry of anti-pH3 (green) labeling cells in M-phase of the cell cycle on 3 dpf frontal retinal cryosections counterstained with the nuclear marker DAPI (blue). *Hdac1*$^{-/-}$ retinae show a higher number of mitotic cells in comparison to wild type retinae at 3 dpf. Scale bar, 50 µm.

**Supplementary Figure 5. *Hdac1* loss-of-function results in cell cycle regulator expression defects in the retina.** Expression analysis of the three cell cycle regulators *c-myc*, *p27* and *cyclinD1* at 2 days post-fertilization (dpf) in wild type (sibling, respectively **A**, **B**, **C**), *hdac1*$^{-/-}$ mutant (**A'**, **B'**, **C'**). (**A – A'**) Comparative *in situ* hybridization of *c-myc* expression reveals an increase in expression of *c-myc* in the central part of *hdac1*$^{-/-}$ mutant retinae in comparison to wild type retinae. (**B – B'**) Frontal view of 2 dpf zebrafish retinae hybridized with *p27* antisense RNA probe. Expression analysis of *p27* reveals a complete loss of *p27* expression in *hdac1*$^{-/-}$ mutant retinae in comparison to wild type retinae. (**C – C'**) Similarly to *c-myc*, comparative analysis of *cyclinD1* expression in wild type (sibling) and *hdac1*$^{-/-}$ mutant retinae shows an increase in the expression of *cyclinD1* in the retinae of *hdac1*$^{-/-}$ mutant in comparison to wild type siblings. Scale bar, 50 µm.

**Supplementary Figure 6. *9*-HSA exogenous administration leads to a decrease in proliferation in the hindbrain.** pH3 staining (brown) labeling mitotic cells of whole-mount embryos injected with DMSO (control, **A**) and *9*-HSA (**B**) at 1 day post-fertilization (dpf). (**C**)

Quantification of the mitotic index in the hindbrain at 1 dpf reveals a significant decrease in the number of proliferative cells in the hindbrain of *9*-HSA -treated compared to DMSO-treated embryos (n = 7 for DMSO and n = 9 for *9*-HSA –treated. Average number of pH3-positive cells in DMSO-treated hindbrain = 246, SD ± 26 and average number of pH3-positive cells in *9*-HSA –treated hindbrain = 203, SD ± 25; *p*-value: 0.005142). ** *p*-value < 0.01. Scale bar, 50 µm.

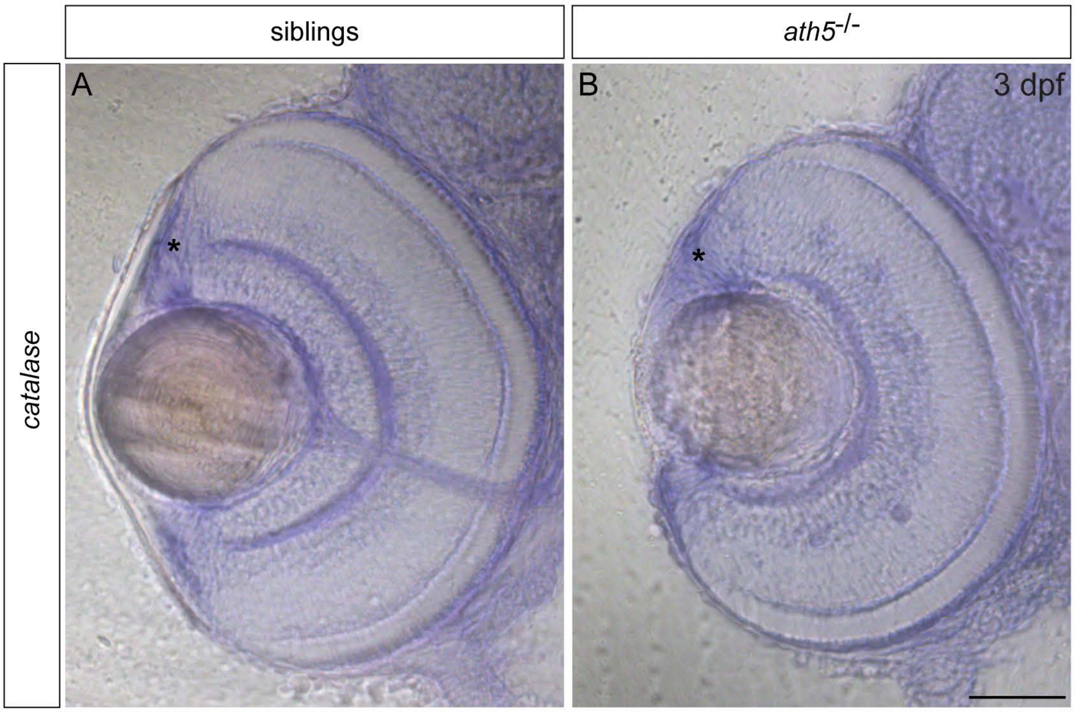


**Supplementary Figure 1**

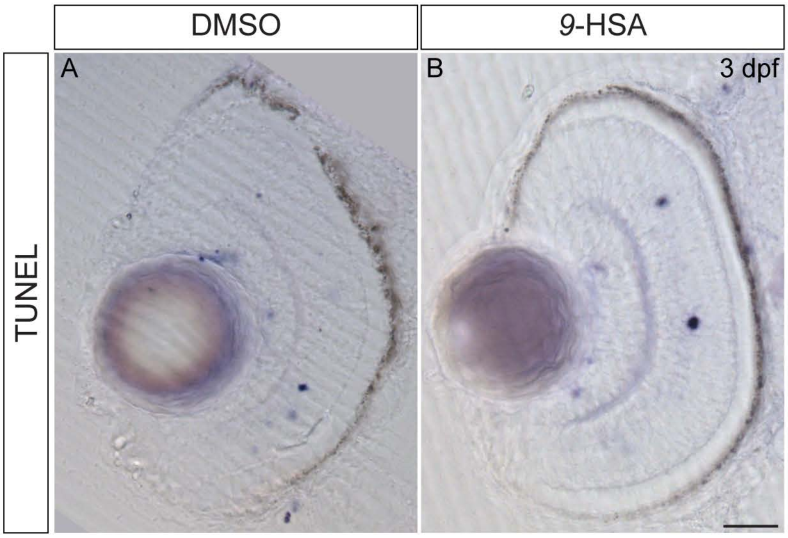


**Supplementary Figure 2**

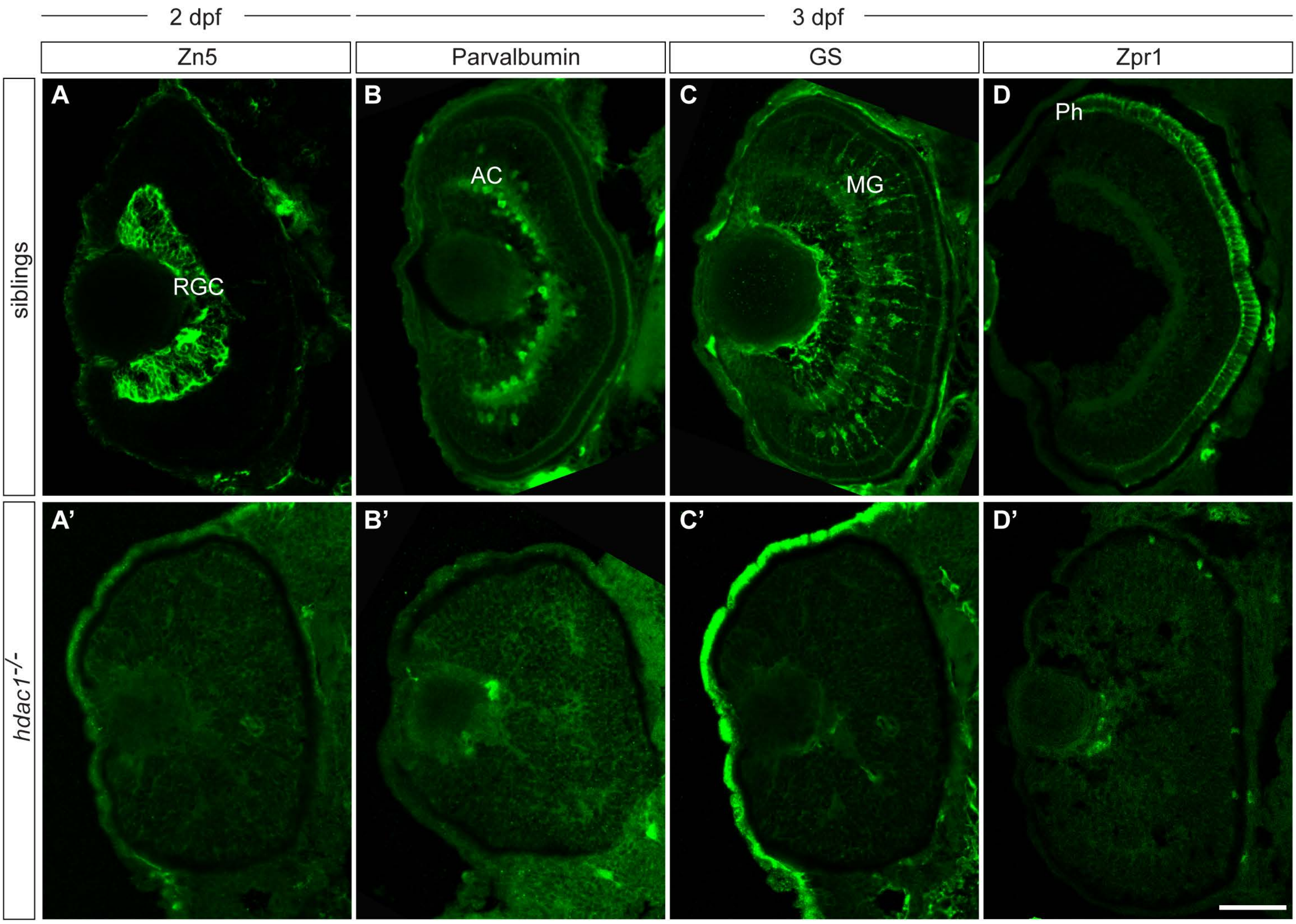


**Supplementary Figure 3**

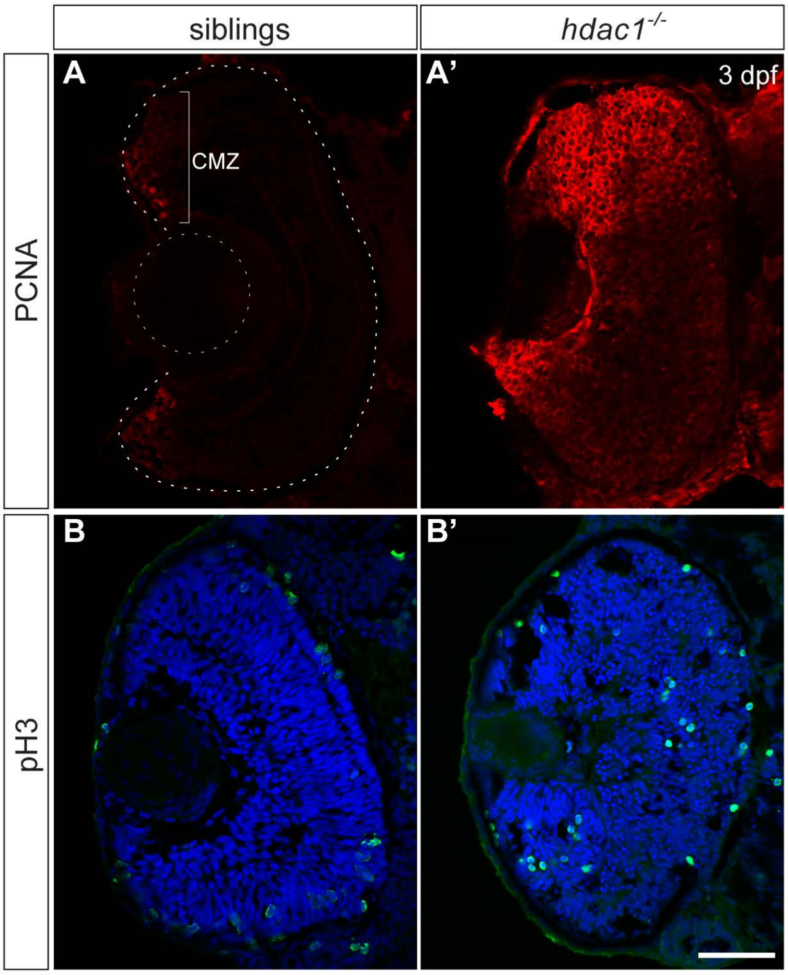


**Supplementary Figure 4**

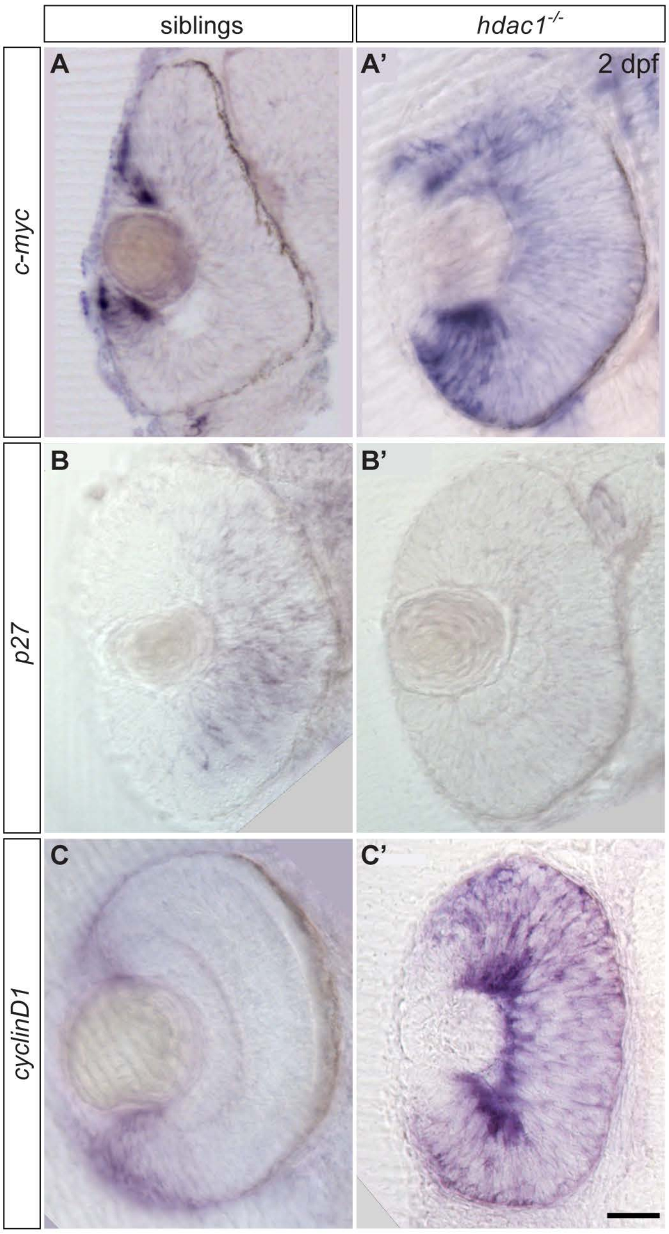


### Supplementary Figure 5

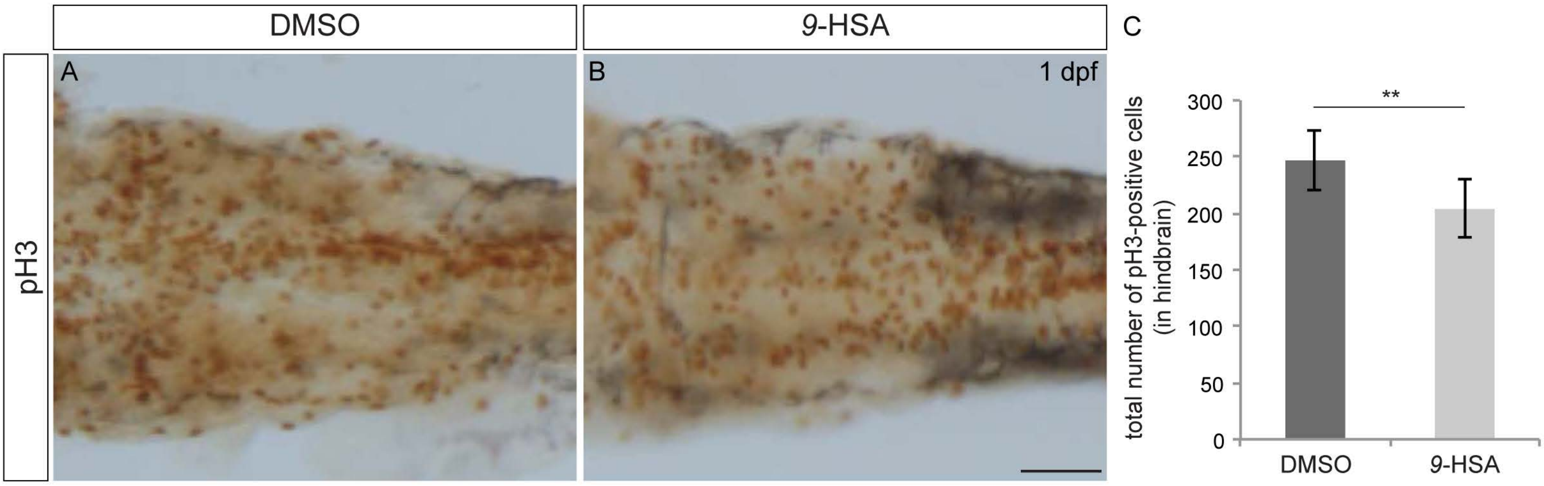


**Supplementary Figure 6**
